# Singlet oxygen-induced signalling depends on the metabolic status of the Chlamydomonas cell

**DOI:** 10.1101/2021.11.03.467076

**Authors:** Waeil Al Youssef, Regina Feil, Maureen Saint-Sorny, Xenie Johnson, John E. Lunn, Bernhard Grimm, Pawel Brzezowski

## Abstract

Using a novel mutant screen, we identified TREHALOSE 6-PHOSPHATE PHOSPHATASE 1 (TSPP1) as a functional enzyme dephosphorylating trehalose 6-phosphate (Tre6P) to trehalose in *Chlamydomonas reinhardtii*. The *tspp1* knock-out results in reprogramming of the cell metabolism via significantly altered transcriptome. As a secondary effect, *tspp1* also shows impairment in ^1^O_2_-induced chloroplast retrograde signalling. From transcriptomic analysis and metabolite profiling, we conclude that accumulation or deficiency of certain metabolites directly affect ^1^O_2_-signalling. ^1^O_2_-inducible *GLUTATHIONE PEROXIDASE 5* (*GPX5*) gene expression is suppressed by increased content of fumarate, an intermediate in the tricarboxylic acid cycle (TCA cycle) in mitochondria and dicarboxylate metabolism in the cytosol, while it is promoted by another TCA cycle intermediate, aconitate. Furthermore, genes encoding known essential components of chloroplast-to-nucleus ^1^O_2_-signalling, PSBP2, MBS, and SAK1, show decreased transcript levels in the *tspp1* mutant, which can be rescued by exogenous application of aconitate. We demonstrate that chloroplast retrograde signalling involving ^1^O_2_ depends on mitochondrial and cytosolic processes and that the metabolic status of the cell determines the response to ^1^O_2_.

## Introduction

Nearly forty years ago, tetrapyrrole biosynthesis (TBS) intermediates were proposed to be involved in chloroplast retrograde (organelle-to-nucleus) signalling. This conclusion was based on the observation that accumulation of chlorophyll precursors negatively affects transcript level of light-harvesting chlorophyll a/b-binding (LHCB) protein^1^. Later studies on chloroplast retrograde signalling involved mutants^2^, treatment with a carotenoid biosynthesis inhibitor, norflurazon^3^, or TBS inhibitors, such as α,α-dipyridyl^1^ or thujaplicin^4^. Experiments with norflurazon led to the discovery of the *<u>g</u>enomes <u>un</u>coupled* (*gun*) mutants with a common phenotype of chloroplast status-independent expression of photosynthesis-associated nuclear genes^3^.

It is noteworthy that treatment with inhibitors causing carotenoid deficiencies results in generation of reactive oxygen species (ROS) and photooxidative damage to chloroplasts. Furthermore, the end-products of TBS, haem and chlorophyll, as well as many of their intermediates (Supplementary Fig. 1) are light-absorbing and redox-reactive molecules, capable to generate ROS. Singlet oxygen (^1^O_2_) can be produced through interaction of ground (triplet)-state oxygen (^3^O_2_) with triplet-state chlorophyll or TBS intermediates, e.g. protoporphyrin IX (Proto), which are excited by light^5^. ROS are also metabolic products of other cellular processes in plants, such as photosynthesis and respiration.

Although ^1^O_2_ is not considered to be the most reactive oxygen species, it is thought to be the major ROS involved in photooxidative damage^6^. However, it was shown that production of ^1^O_2_ in the chloroplast induces stress responses that do not result exclusively from physicochemical damage, but also rely on signal transduction triggered by ROS^7^. ^1^O_2_ has a short half-life (about 200 ns) in the cell^8^ and as a result, the distance that it may move was calculated to be approximately 10 nm, based on predicted diffusion rates^9,10^. Its diffusion range is also limited due to its high reactivity with membrane lipids^11^. Therefore, ^1^O_2_ could play a specific role as an activator of a stress response only if it is detected close to its source, which strongly suggests that other components mediating the ^1^O_2_ signals should exist. Alternatively, altered metabolite contents triggered by ^1^O_2_ may also mediate ^1^O_2_-retrograde signalling, or certain metabolic signature may be required to trigger changes in nuclear gene expression^12,13^.

Despite the profound effect of ^1^O_2_ on the chloroplast redox state and its apparent involvement in altering nuclear gene expression, little is known about the components involved in ^1^O_2_-dependent retrograde signalling. The protein factors identified so far include EXECUTER1 (EX1) and EX2 in the *fluorescence* (*flu*) mutant of *A. thaliana*^14^, the P-subunit of photosystem II family protein (PSBP2)^15^ and SINGLET OXYGEN ACCLIMATION KNOCKEDOUT1 (SAK1) in *Chlamydomonas reinhardtii*^16^, or METHYLENE BLUE SENSITIVITY (MBS1) shown to be involved in ^1^O_2_-signalling in both *A. thaliana* and *C. reinhardtii*^17^.

We hypothesized that ^1^O_2_ generated via the photosensitizing activity of Proto, rather than porphyrin accumulation itself, triggers signalling cascades that alter nuclear gene expression. Therefore, we used a *C. reinhardtii* mutant *chlD-1*^18^ that does not produce chlorophyll and accumulates Proto due to a dysfunctional Mg-chelatase (MgCh, Supplementary Fig. 1), but with introduced over-expression of GUN4, *chlD-1/GUN4*^19^. Endogenous accumulation of Proto in mutants is advantageous for studying ^1^O_2_-signalling, because it eliminates the need for exogenous application of TBS or carotenoid biosynthesis inhibitors, as well as photosensitizers, such as rose bengal or neutral red to induce ^1^O_2_ generation^15,20,21^, which are not natural products of the cell, do not localize specifically to any subcellular compartment, and as a consequence may result in artefactual responses. The lack of chlorophyll provides another advantage, because such mutants do not have functional photosynthetic electron transport, so that ^1^O_2_ production and signaling originating in photosynthesis is avoided. The accumulating Proto is thus the dominant source of generated ^1^O_2_ in the chloroplast and we hypothesized that this approach should allow us to isolate novel components or mechanisms governing ^1^O_2_-signaling. The GENOMES UNCOUPLED 4 (GUN4) protein involved in MgCh function (Supplementary Fig. 1) and signalling degrades upon Proto accumulation^19^. However, *chlD-1* overexpressing GUN4 showed higher GUN4 content than *chlD-1*, while it retained the chlorophyll-free phenotype^19^. Additionally, *chlD-1/GUN4* demonstrated higher expression of *GLUTATHIONE PEROXIDASE 5* (*GPX5*) and higher tolerance to ^1^O_2_ than *chlD-1*^19^. Thus, to minimize the GUN4-deficient phenotype and to maintain high and stable ^1^O_2_-inducibility of *GPX5* expression, *chlD-1/GUN4* instead of *chlD-1* was used as the receiver strain for the gene construct that allowed us to monitor ^1^O_2_-signalling. The gene construct introduced into the *chlD-1/GUN4* genome consisted of the promoter region of the *GPX5* gene fused to the promoterless *ARYLSULFATASE 2* gene (*ARS2*). *GPX5* was shown to be specifically induced by ^1^O_2_ in *C. reinhardtii*^20,22^, while the ARS2 activity can be assessed by the enzymatic assay^23^. The resulting *GPX5-ARS2* gene construct was shown previously to be an effective reporter to study ^1^O_2_-signalling^15,22,24^. Following introduction of *GPX5-ARS2*, transformant strain showing high inducibility of *GPX5-ARS2* in response to ^1^O_2_ was named *<u>sig</u>nalling <u>Rep</u>orter* (*sigRep*) and was subjected to further studies.

Subsequent random insertional mutagenesis of *sigRep*, followed by a screening for decreased *GPX5-ARS2* expression identified nine mutants with impaired ^1^O_2_-signalling. To reflect the impairment in ^1^O_2_-dependent signalling, these mutants were named *<u>g</u>enomes <u>un</u>coupled <u>S</u>inglet <u>O</u>xygen <u>S</u>ignalling* (*gunSOS*). The *gunSOS1* mutant was selected for further analysis. Although employment of the Proto-accumulating mutant as the background strain provides advantage in studying ^1^O_2_-signalling, it also results in certain limitations, e.g. lack of chlorophyll enforces heterotrophic growth, alters nuclear gene expression and metabolism. Thus, due to the character of the background strain, phenotype of *gunSOS1* was primarily compared to the parental *sigRep*, instead of the chlorophyll-synthesizing wild type (WT).

The lack of the retrograde response to ^1^O_2_ in *gunSOS1* was verified by decreased expression of *GPX5*, as well as *SAK1, MBS*, and *PSBP2* compared to *sigRep*. The causal mutation in *gunSOS1* was found in a gene annotated as *TREHALOSE 6-PHOSPHATE PHOSPHATASE* (hereafter *TSPP1*). Besides clear impairment in ^1^O_2_-signalling, mutation in *TSPP1* also resulted in accumulation of trehalose 6-phosphate (Tre6P) in *gunSOS1* compared to *sigRep* and WT. Therefore, TSPP1 is the first confirmed phosphatase acting on Tre6P in *C. reinhardtii* and in fact the first enzyme with a confirmed function in trehalose metabolism in this organism. Our data indicate that accumulation of Tre6P, alternatively the lack of the TSPP1 protein, causes changes to the expression of genes involved in several metabolic pathways. However, Tre6P or TSPP1 are rather intermediates than the primary cause of the impaired ^1^O_2_-signalling. Here, using a combination of genetics, gene expression and metabolic phenotypes, we reveal a complex interaction between chloroplast, mitochondria and cytosol in ^1^O_2_ signal transmission to the nucleus.

## Results

### Forward genetics screen to identify components of the ^1^O_2_-signalling

To generate a mutant impaired in ^1^O_2_-dependent signalling, we first created a reporter strain in a known ^1^O_2_-generating mutant, which was subsequently subjected to mutagenesis. The *GPX5-ARS2* reporter construct (Supplementary Fig. 2) was introduced into the genome of *chlD-1/GUN4* to express the ARS2 protein in a ^1^O_2_-dependent manner, and the transformants were tested for the enzymatic activity of ARS2. The reporter strain, which showed the lowest ARS2 activity in darkness and the highest activity in light was named *<u>sig</u>nalling <u>Rep</u>orter* (*sigRep*; Fig. 1A) and was used in further applications. Expression kinetics of the cytosolic (*GPX5_cyt_*) and chloroplastic (*GPX5_cp_*) version of *GPX5*^25^; Fig. 1b), as well as the *GPX5-ARS2* construct (Fig. 1c) were examined by quantitative Real-Time PCR (qRT-PCR) upon transfer from dark to light. As expected, higher induction of *GPX5_cp_* and *GPX5_cyt_* were observed upon light illumination in *sigRep* compared to WT (Fig. 1b). Induced *GPX5-ARS2* expression in *sigRep* in the light (Fig. 1c) confirmed the results obtained in the ARS activity assay (Fig. 1a) and a suitability of the *sigRep* reporter strain for further applications.

**Fig. 1.**
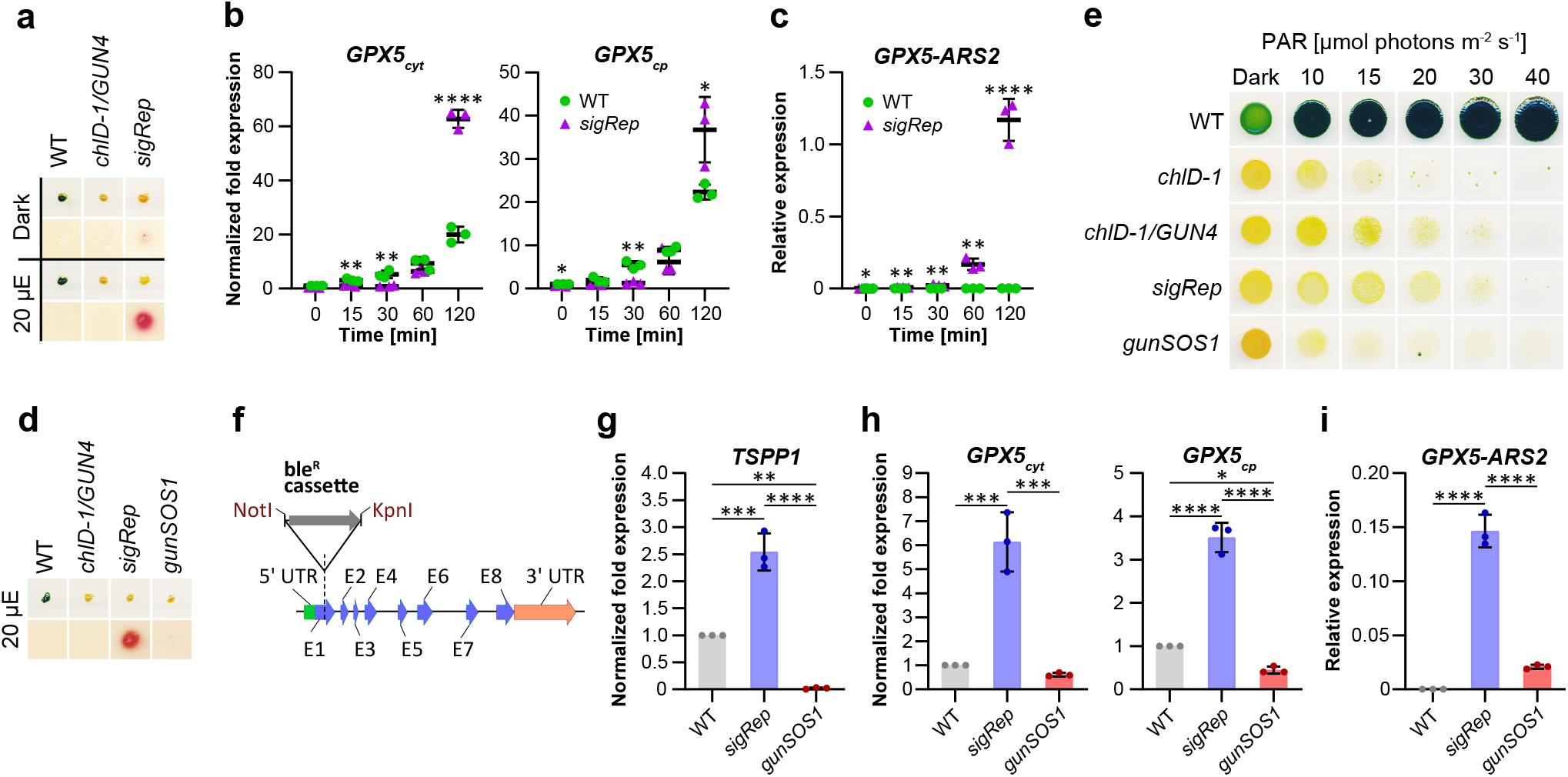
Generation and initial characterization of the mutant impaired in ^1^O_2_-signalling. **a** Arylsulfatase (ARS2) assay for selection of the strain expressing *GPX5-ARS2* reporter gene in a ^*1*^O_2_-inducible manner. The mutant selected for further applications was named *<u>sig</u>nalling <u>Rep</u>orter* (*sigRep*). WT and *chlD-1/GUN4* do not carry *GPX5-ARS2* and were used as negative controls in ARS2 assay. **b** Expression kinetics of cytosolic (*GPX5_cyt_*) and chloroplast *GPX5* (*GPX5_cp_*) in *sigRep* compared to WT upon exposure to light; the values were normalized to WT in dark. **c** Expression of *GPX5-ARS2* in *sigRep* positively correlates with time of exposure to light. WT was used as a negative control. **d** ARS2 activity assay following mutagenesis of *sigRep*. Screen was performed to identify mutants not expressing *GPX5-ARS2* compared to *sigRep* in the same conditions. The mutant selected for further applications was named *genomes <u>un</u>coupled <u>S</u>inglet <u>O</u>xygen <u>S</u>ignalling <u>1</u>* (*gunSOS1*). WT and *chlD-1/GUN4* were used as negative controls. **e** Strains with impaired ^1^O_2_-signalling demonstrated increased sensitivity to light compared to *sigRep* and *chlD-1/GUN4*. Higher light tolerance of *chlD-1/GUN4* and *sigRep* compared to *chlD-1* is also visible. **f** *TSPP1* (Cre12.g497750) gene model. The insertion (ble^R^) was identified in the first exon of 3241 bp *TSPP1*. **g** Expression of *TSPP1* in *gunSOS1* compared to *sigRep*. **h** Expression of *GPX5_cyt_* and *GPX5_cp_* in *gunSOS1* compared to *sigRep*. **i** Expression of *GPX5-ARS2* in *gunSOS1* compared to *sigRep*. WT was used as a negative control. Transcript analyses were performed by qRT-PCR on biological triplicates, values were calculated either as a normalized fold expression (2^−ΔΔCt^; WT = 1) or relative expression (2^ΔCt^). For (b) and (c), horizontal bars represent the calculated mean (*n* = 3), vertical error bars represent calculated ±SD; significant differences were calculated using two-tailed Student’s *t*-test and are indicated by asterisks (non-significant not shown), \**P* < 0.05, \*\**P* < 0.01, \*\*\**P* < 0.001, and ****P < 0.0001. For (g)-(i), error bars indicate calculated ±SD; one-way ANOVA, pair-wise comparison with the Tukey’s post-hoc test (non-significant not shown), \**P* < 0.05, \*\**P* < 0.01, \*\*\**P* < 0.001, and \*\*\*\**P* < 0.0001.

Subsequently, a random mutagenesis was performed on *sigRep* using bleomycin resistance cassette (ble^R^) as an insert, followed by screening to isolate strains with lower or undetectable ARS2 activity in the light compared to *sigRep*, and thus, possibly disrupted ^1^O_2_-dependent signalling. Screen of 804 transformants allowed us to isolate nine, which were named *<u>g</u>enomes <u>un</u>coupled <u>S</u>inglet <u>O</u>xygen <u>S</u>ignalling* (*gunSOS*). The *gunSOS1* mutant (Fig. 1d) was selected for further analysis. Higher sensitivity to light was recorded in *gunSOS1* compared to the *sigRep* background strain (Fig. 1e), in agreement with a higher sensitivity or impaired acclimation to ^1^O_2_. Mutant impaired in ^1^O_2_-signalling showed similar Proto accumulation in light compared to *sigRep* and *chlD-1/GUN4* (Supplementary Fig. 3), which eliminated the possibility that spurious mutations in *gunSOS1*, which may have been introduced during mutagenesis, resulted in reduced content of this photosensitizer. Because Proto is considered as the main source of ^1^O_2_, a lower levels of this ROS in *gunSOS1* compared to *sigRep* would not be expected.

The ble^R^ insertion in *gunSOS1* was located in the first exon of the gene annotated as *TREHALOSE 6-PHOSPHATE PHOSPHATASE 1* (here *TSPP1*; locus Cre12.g497750) in the JGI portal (Department of Energy Joint Genome Institute, https://phytozome-next.jgi.doe.gov, v13, genome v5.6; Fig. 1f). The *TSPP1* transcript abundance was determined in *gunSOS1* and compared to *sigRep* by qRT-PCR using primers annealing to the coding sequence upstream of the insertion site. At 2 h after transfer from dark to light, *TSPP1* mRNA content increased 2.5-fold in *sigRep* compared to WT, while it was nearly absent in *gunSOS1* (Fig. 1g). Expression of *GPX5_cyt_* and *GPX5_cp_* was about 10 and 8 times lower in *gunSOS1* compared to *sigRep*, respectively (Fig. 1h). The transcript abundance for *GPX5-ARS2* was 7 times lower in *gunSOS1* than in *sigRep* (Fig. 1i), and explains undetectable ARS2 activity in the ^1^O_2_-signalling mutant (Fig. 1d).

### *C. reinhardtii* TSPP1 is a functional Tre6P phosphatase

While *gunSOS1* is clearly impaired in ^1^O_2_-signalling, it was necessary to determine the primary phenotype caused by the mutation in *TSPP1*. The content of Tre6P in *sigRep* and *gunSOS1* was determined by anion-exchange high performance liquid chromatography coupled to tandem mass spectrometry (LC-MS/MS). Tre6P accumulation increased in *gunSOS1* upon exposure to light in a time-dependent manner, while it decreased in *sigRep* (Fig. 2a). Accumulation of Tre6P in *gunSOS1* indicates that *C. reinhardtii* TSPP1 is a functional phosphatase dephosphorylating Tre6P to trehalose. Subsequently, *TSPP1* expression was found to be induced in the light in *sigRep*, but not in the WT, indicating its inducibility by photooxidative stress rather than light (Fig. 2b). Mutation of the *TSPP1* gene would be expected to decrease dephosphorylation of Tre6P and lower trehalose content, but we also observed accumulation of trehalose in *gunSOS1* (Fig. 2c). This apparent discrepancy can be explained by the existence of another enzyme in *C. reinhardtii* with possible Tre6P phosphatase activity, in combination with higher levels of Tre6P (see Discussion section).

**Fig. 2.**
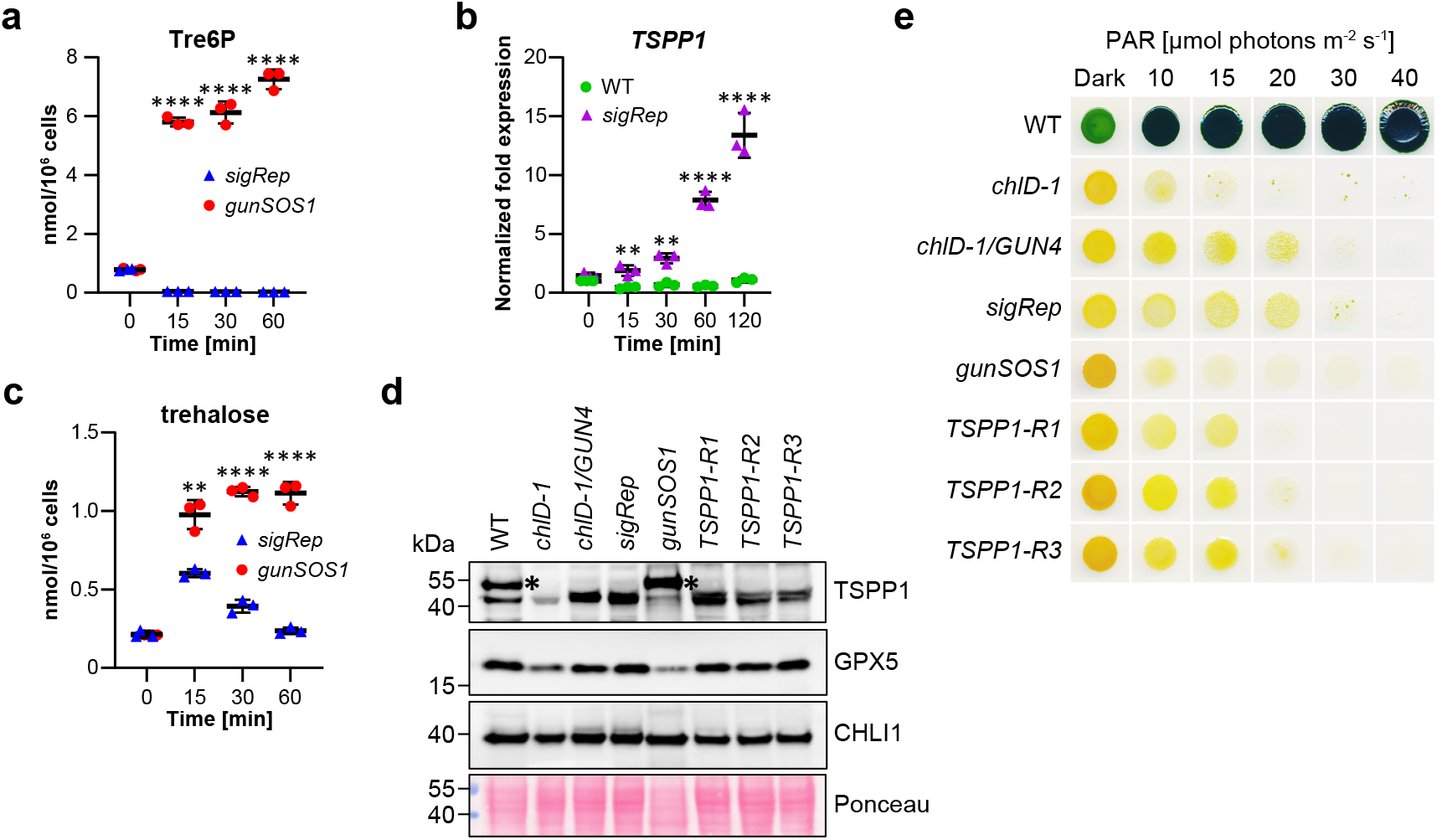
*C. reinhardtii* TSPP1 is a functional phosphatase induced during photooxidative stress. **a** Kinetics of Tre6P accumulation in *gunSOS1* compared to *sigRep* upon transfer from dark to light. **b** Kinetics of *TSPP1* expression upon transfer from dark to light in *gunSOS1* compared to WT indicate inducibility by photooxidative stress rather than light; values were calculated as a normalized fold expression (2^−ΔΔCt^; WT = 1). **c** Higher content of trehalose in *gunSOS1* compared to *sigRep* upon transfer from dark to light may be resulting from the possible phosphatase activity of TSSP1, which was not determined in the present study. **d** Immuno-blot analysis of TSPP1 (calculated MW of 42 kDa) and GPX5 content in *gunSOS1* compared to *sigRep* and strains rescued with wild-type copy of *TSPP1*. The CHLI1 content was used as a loading control. The unspecific immunoreaction is indicated by an asterisk. **e** Light sensitivity examination showed that the strains with rescued ^1^O_2_-signalling have an increased tolerance to light compared to *gunSOS1*. Experiments in (a), (b) and (c) were performed in biological triplicates (*n* = 3), horizontal bars represent the calculated mean, vertical error bars represent ±SD; significant differences were calculated using two-tailed Student’s *t*-test and are indicated by asterisks (non-significant not shown), \**P* < 0.05, \*\**P* < 0.01, \*\*\**P* < 0.001, and \*\*\*\**P* < 0.0001.

Rescue of the TSPP1 deficiency in *gunSOS1* was performed with the isolated genomic DNA fragment carrying the wild-type *TSPP1* gene. Several independent transformants showed rescued ^1^O_2_-signalling, which was indicated by the *GPX5-ARS2* expression determined in the ARS assay, and data from three representative strains are shown in Supplementary Fig. 4a. The analysed rescued strains also showed increased *GPX5_cp_* transcript content compared to *gunSOS1* (Supplementary Fig. 4b). A peptide-specific antibody for TSPP1 was produced to compare the protein content in *gunSOS1* to strains showing ^1^O_2_-dependent signalling. A faint immune signal of approximately 43 kDa was detected in *gunSOS1* and *chlD-1*, but two narrow bands were seen in *chlD-1/GUN4, sigRep* and the *TSPP1*-rescued strains (Fig. 2d). WT and *gunSOS1* showed an additional unspecific immunoreaction with protein of approximately 53 kDa. As expected, GPX5 content was lower in *gunSOS1* compared to *sigRep* (Fig. 2d), which correlated with qRT-PCR results on *GPX5* transcript levels (Fig. 1h). Strains with rescued ^1^O_2_-signalling, *TSPP1-R1, -R2, and -R3*, showed increased content of GPX5 (Fig. 2d) and higher tolerance to light compared to *gunSOS1* (Fig. 2e).

### The *gunSOS1* mutant shows altered metabolism compared to *sigRep*

In *A. thaliana* and other angiosperms, Tre6P functions as a sucrose signal and homeostatic regulator of sucrose metabolism^26,27^, which links plant growth and development to the availability of sucrose^28^, reviewed in^29^. To determine the effect of accumulated Tre6P on metabolism in *C. reinhardtii*, we performed comparative metabolite profiling of *gunSOS1* and *sigRep* (Supplementary Table 1), which revealed a relatively wide range of variation. Aside from Tre6P and trehalose, 11 out of 27 analysed metabolites showed significantly increased content in *gunSOS1* compared to *sigRep* in the light (Supplementary Fig. 5a). By means of the MetaboAnalyst 5.0 portal (https://www.metaboanalyst.ca) these metabolites were assigned to the metabolic processes in the cell. A hypergeometric test indicated the most affected metabolic pathways, i.e. the TCA cycle, pyruvate metabolism, and starch and sucrose metabolism (Supplementary Fig. 5b and Supplementary Table 2). Eight metabolites showed significantly decreased content in *gunSOS1* compared to *sigRep* (Supplementary Fig. 6a), with the highest impact (MetaboAnalyst 5.0) on fructose and mannose metabolism, amino sugar and nucleotide sugar metabolism, and starch and sucrose metabolism (Supplementary Fig. 6b and Supplementary Table 3). Metabolites, which did not show a significant change in *gunSOS1* compared to *sigRep* are presented in Supplementary Fig. 7. The map-overview of the selected metabolic pathways with depicted metabolites that showed different content, or were not changed, in *gunSOS1* relative to *sigRep* is presented in Fig. 3.

**Fig. 3.**
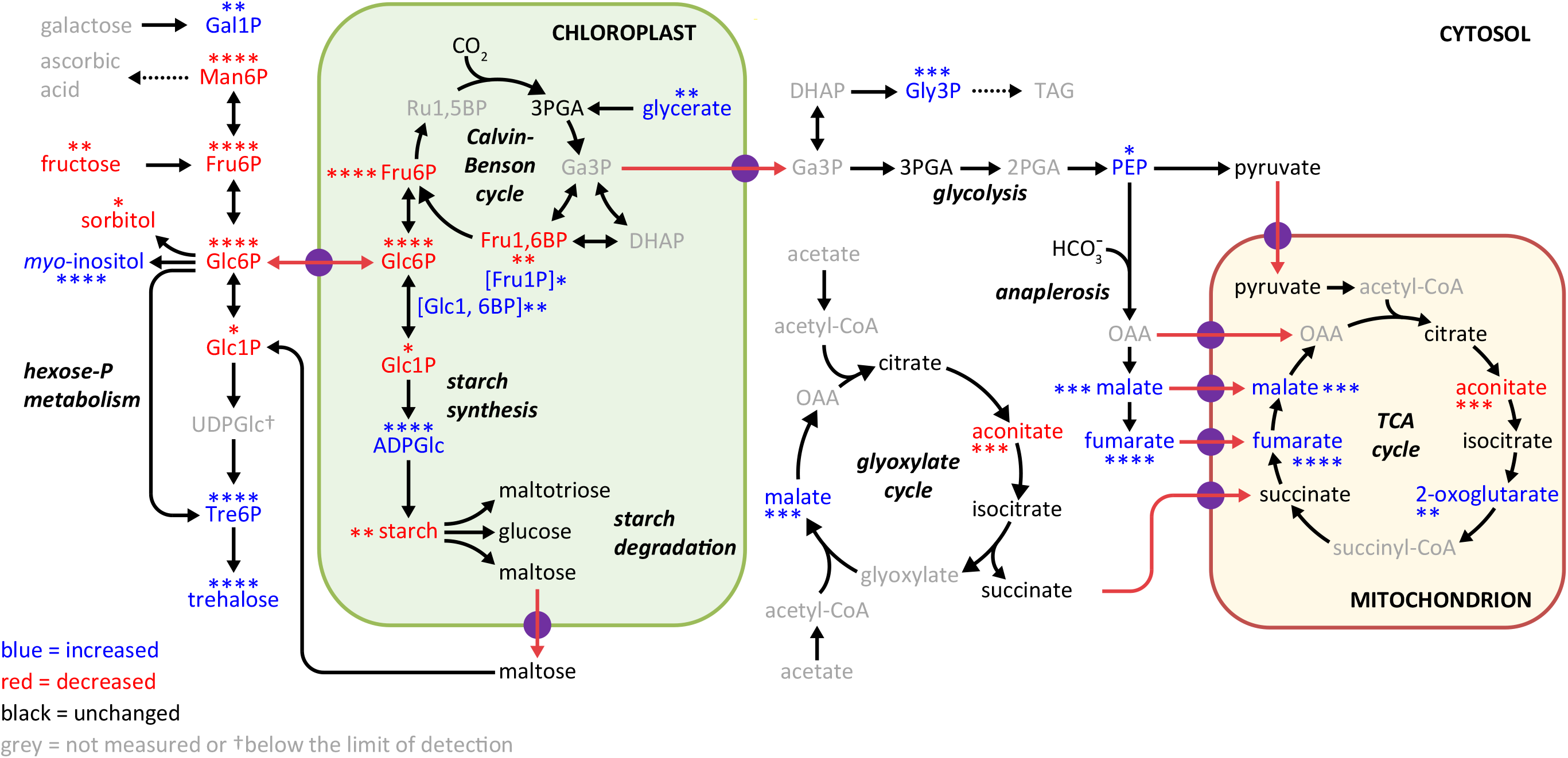
The overview of the selected metabolic pathways in *gunSOS1* compared to *sigRep*. The map was constructed based on the measured metabolites presented in Supplementary Figures 5a, 6a, and 7. Metabolites increased in *gunSOS1* compared to *sigRep*: fumaric acid (fumarate), oxoglutaric acid (2-OG), myo-inositol, ADP-glucose (ADPGlc), galactose 1-phosphate (Gal1P), fructose 1-phosphate (Fru1P), alpha-D-glucose 1,6-bisphosphate (Glc1, 6BP), glycerol 3-phosphate (Gly3P), phosphoenolpyruvic acid (PEP), L-malic acid (malate). Metabolites decreased in *gunSOS1* compared to *sigRep*: cis-aconitic acid (aconitate), mannose 6-phosphate (Man6P), glucose 6-phosphate (Glc6P), D-fructose (fructose), fructose 6-phosphate (Fru6P), glucose 1-phosphate (Glc1P), fructose 1,6-bisphosphate (Fru1, 6BP), and sorbitol. Metabolites that did not show significant difference in *gunSOS1* relative to *sigRep*: D-glucose (glucose), D-maltose (maltose), maltotriose, citric acid (citrate), isocitric acid (isocitrate), pyruvic acid (pyruvate), succinic acid (succinate), 3-phosphoglyceric acid (3PGA). The colour-coding is shown in the legend. Measurements were performed in biological triplicates (*n* = 3); significant differences were calculated using two-tailed Student’s *t*-test. The maximal significant differences from any time point in light (Supplementary Figures 5a and 6a) are indicated by asterisks, \**P* < 0.05, \*\**P* < 0.01, \*\*\**P* < 0.001, and ****P < 0.0001. The metabolic map was constructed based on the review by Johnson and Alric^103^ and Findinier et al.^104^.

Starch content was determined in *gunSOS1* and compared to *sigRep*, because of changes in several intermediates of starch and sucrose metabolism (Supplementary Figs. 5b and 6b). Indeed, *gunSOS1* accumulated only 60% of the starch content of *sigRep* when the strains were grown in acetate-supplemented media (TAP, Supplementary Fig. 6c). Both *sigRep* and *gunSOS1* are obligate heterotrophs. *SigRep* contained less starch after transfer from TAP and cultivation in TP for 48 h, whereas the starch content of *gunSOS1* was essentially unchanged (Supplementary Fig. 6c), indicating that there was net degradation of starch in *sigRep* but not in *gunSOS1*. Starch formation is strongly induced during nitrogen (N) deprivation^30^. The *sigRep* cells did not increase starch reserves after 3 days in N-deficient medium, suggesting that starch accumulation had already reached its maximum under non-stress conditions. The starch content in *gunSOS1* deprived of N did increase by 50% and was similar to the content in *sigRep* (Supplementary Fig. 6c). We conclude that *sigRep* accumulates high levels of starch as a consequence of the *chlD-1/GUN4* genetic background and that the accumulation of Tre6P represses this response in *gunSOS1* leading to a lower accumulation of starch. The *gunSOS1* cells are still capable of responding to the N-deprivation stress and accumulating starch despite disturbed metabolism. Starch degradation on the other hand, is compromised in *gunSOS1*, which could indicate a general impairment of catabolic processes or a defect in sensing carbon limitation.

### Increased Tre6P content alters metabolism in *C. reinhardtii* via transcriptional changes

To determine the extent to which chloroplast retrograde signalling is affected in *gunSOS1*, we performed comparative RNA-sequencing (RNA-seq) of the transcriptomes of *gunSOS1* and *sigRep* upon transfer from dark to light. The analysis showed that 8120 genes were expressed in both strains, but 1377 and 1087 additional genes were uniquely expressed in *gunSOS1* and *sigRep*, respectively (Fig. 4a, detailed listing can be found in Supplementary Dataset 1). Furthermore, differential gene expression analysis showed that the transcript contents of 1606 genes and 1706 genes were lower or higher, respectively, in *gunSOS1* compared to *sigRep* (Supplementary Dataset 2).

**Fig. 4.**
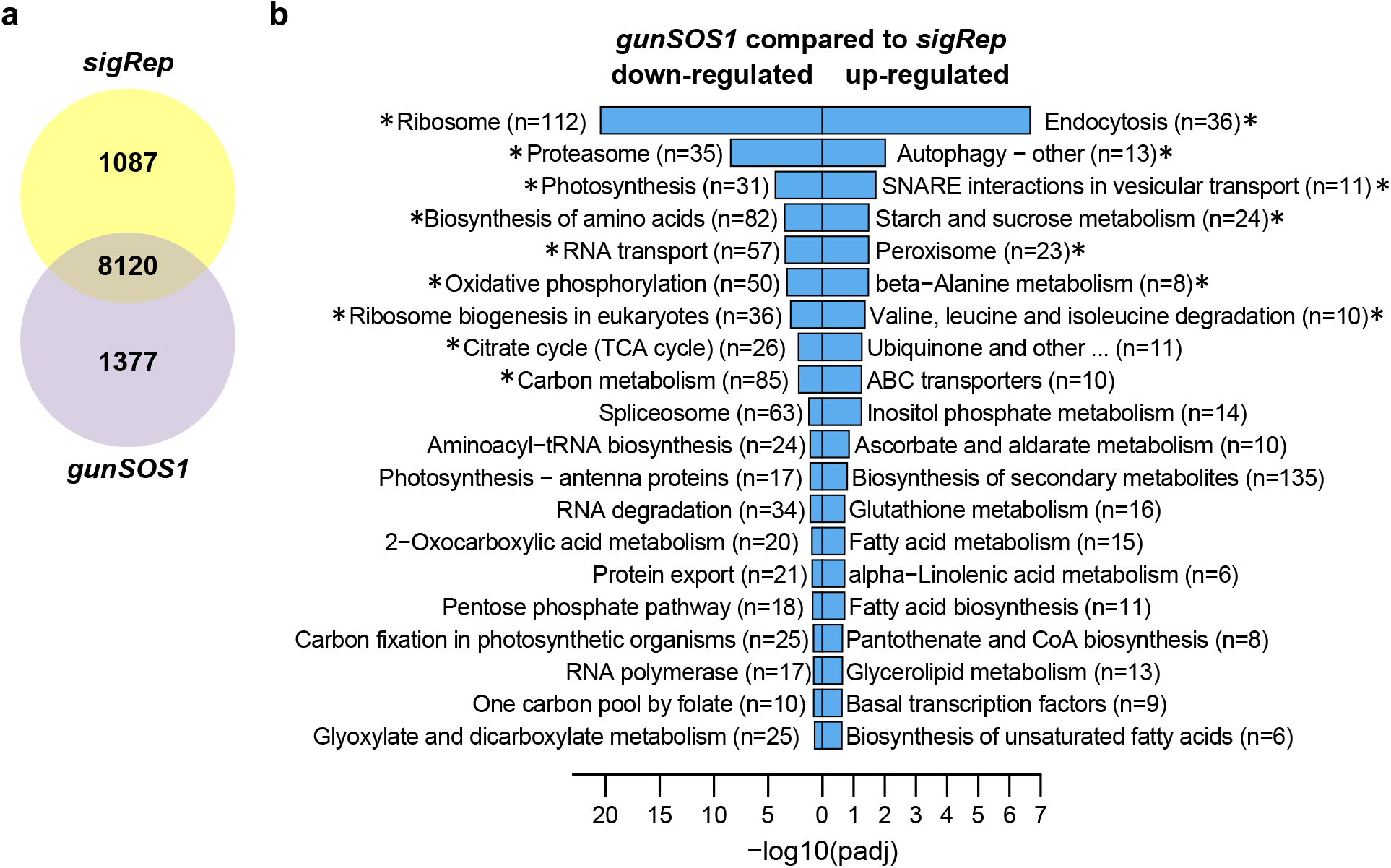
Transcriptome analysis based on RNA-seq. **a** Venn diagram showing number of genes expressed in *gunSOS1* and *sigRep* (8120), or uniquely expressed in *gunSOS1* (1377) or in *sigRep* (1087). **b** Pathway enrichment analysis (KEGG, http://www.kegg.jp/) based on the differentially expressed genes (RNA-seq). Effect on metabolic processes is sorted by the *P* value. Asterisks in (b) indicate the core (significant) enrichment.

Based on the KEGG ontology (http://www.kegg.jp/) several genes associated with metabolic pathways were down- or up-regulated (Fig. 4b) in *gunSOS1* compared to *sigRep* (Supplementary Dataset 3). Given the strong overlap between differentially expressed genes and corresponding pathway intermediates, we hypothesized that the altered metabolism can be directly explained by altered gene expression. To decipher a complex correlation between altered signalling, gene expression and metabolism, accumulation or deficiency in metabolites from given pathway(s) was correlated with altered gene expression in *gunSOS1* compared to *sigRep*. Fumarate and aconitate will be described in more detail, as representative examples of the metabolites showing respectively increased or decreased content in *gunSOS1* compared to *sigRep*.

KEGG enrichment of the RNA-seq data indicated increased expression of the gene encoding FUMARATE HYDRATASE CLASS II (FUM2, Enzyme Commission (EC) Number 4.2.1.2, Cre01.g020223) in *gunSOS1* compared to *sigRep*. FUM is responsible for reversible stereospecific interconversion of malate to fumarate. However, specific isoforms of this protein act in defined pathways and may favour one direction over the other depending on the subcellular localisation and environment^31^. The mitochondrial isoform catalyses hydration of fumarate to L-malate in the TCA cycle, while the cytosolic form catalyses dehydration of L-malate to fumarate ^32,33^. Two isoforms exist in *C. reinhardtii*, FUM1 encoded in locus Cre06.g254400 and FUM2. Based on the predicted subcellular localisation (PredAlgo; http://lobosphaera.ibpc.fr/cgi-bin/predalgodb2.perl?page=main), FUM1 localizes to the mitochondria, while FUM2 is assigned to the “other” compartment and is likely to be a cytosolic protein. Pathway (KEGG) diagrams for the TCA cycle and pyruvate metabolism can be found in Supplementary Fig. 8, showing integration of metabolite, RNA-seq and qRT-PCR data. Comparison of the transcriptomes of *gunSOS1* and *sigRep* also indicated increased expression of the *FUMARYLACETOACETASE* gene (locus *Cre17.g732802*, EC 3.7.1.2) involved in tyrosine catabolism. FUMARYLACETOACETASE catalyses hydrolysis of 4-fumarylacetoacetate to acetoacetate and fumarate (Supplementary Fig. 9). Increased expression of genes encoding FUM2 and FUMARYLACETOACETASE could explain accumulation of fumarate in *gunSOS1* compared to *sigRep*.

Aconitate deficiency in *gunSOS1* compared to *sigRep* can be explained by transcriptional downregulation of *ACONITATE HYDRATASE (ACH1*, Cre01.g042750, EC 4.2.1.3). ACH1 interconverts citrate and isocitrate, via cis-aconitate, in the TCA cycle (Supplementary Fig. 10) and is also involved in glyoxylate and dicarboxylate metabolism. At equilibrium, the reactants of the ACH1 reaction are present in the following ratio: 91% citrate, 6% isocitrate and 3% aconitate. With the smallest pool of the three tricarboxylic acids, fluctuations in the level of aconitate are expected to be more pronounced compared to the other metabolites involved, especially citrate, whose content was similar in *gunSOS1* and *sigRep* in the light (Supplementary Fig. 7). Taken together, our data indicate that altered fumarate and aconitate content in *gunSOS1* compared to *sigRep* can be explained by altered expression of genes encoding enzymes involved in key metabolic processes affecting those metabolites.

### Impaired O_2_-signalling in *gunSOS1* correlates with decreased expression of *PSBP2, MBS*, and *SAK1*

The PSBP2^15^, MBS^17^, and SAK1^16^ proteins are required for ^1^O_2_-induced chloroplast retrograde signalling in *C. reinhardtii*. Therefore, we determined the transcript levels of *PSBP2, MBS*, and *SAK1*, which were 8-, 9-, and nearly 41-fold lower, respectively, in *gunSOS1* compared to *sigRep* (Fig. 5a). Decreased expression of *SAK1* in *gunSOS1* prompted us to compare transcript levels of selected ^1^O_2_-responsive genes between *gunSOS1* and the *sak1* mutant, presented in Wakao et al.^16^. SAK1 is a key regulator of the gene expression response and its knockout abolishes acclimation response to ^1^O_2_^16^. The involvement of SAK1 in ^1^O_2_-signalling was determined following treatment with rose bengal^16^, which produces ^1^O_2_ in the light, while in *gunSOS1* the main source of ^1^O_2_ is endogenously accumulating Proto (Supplementary Fig. 3). Nevertheless, based on our qRT-PCR analysis, all tested genes had the same reduced inducibility in *gunSOS1* compared to *sigRep* (Fig. 5b), as it was observed in *sak1* compared to its corresponding WT following ^1^O_2_-exposure^16^. Among the genes that were down-regulated in both *gunSOS1* (Fig. 5b) and *sak1*^16^ were two *CYCLOPROPANE FATTY ACID SYNTHASES, CFA1* and *CFA2* (*CPLD27*), involved in lipid and sterol metabolism, as well as the gene encoding SOUL1 heme-binding protein (Fig. 5b). It is noteworthy that attenuated expression of both *CFA1* and *CFA2*, as well as *SOUL1*, was also observed in studies on a *gpx5* mutant^34^. Furthermore, in agreement with the *sak1* phenotype presented in Wakao, et al. ^16^, *gunSOS1* also showed decreases in transcript content of *PEPTIDE METHIONINE SULFOXIDE REDUCTASE* (*MSRA3*), *ALCOHOL DEHYDROGENASE* (*ADH7*), *RETINALDEHYDE BINDING PROTEIN-RELATED* (*RABPR2*), and Cre01.g007300 encoding N-ACETYLPHOSPHATIDYLETHANOLAMINE-HYDROLYZING PHOSPHOLIPASE D (Fig. 5b). Thus, there is an overlap between the phenotypes of *gunSOS1* and *sak1*^16^ with respect to their attenuated expression of genes induced during elevated ^1^O_2_ (Fig. 5b).

**Fig. 5.**
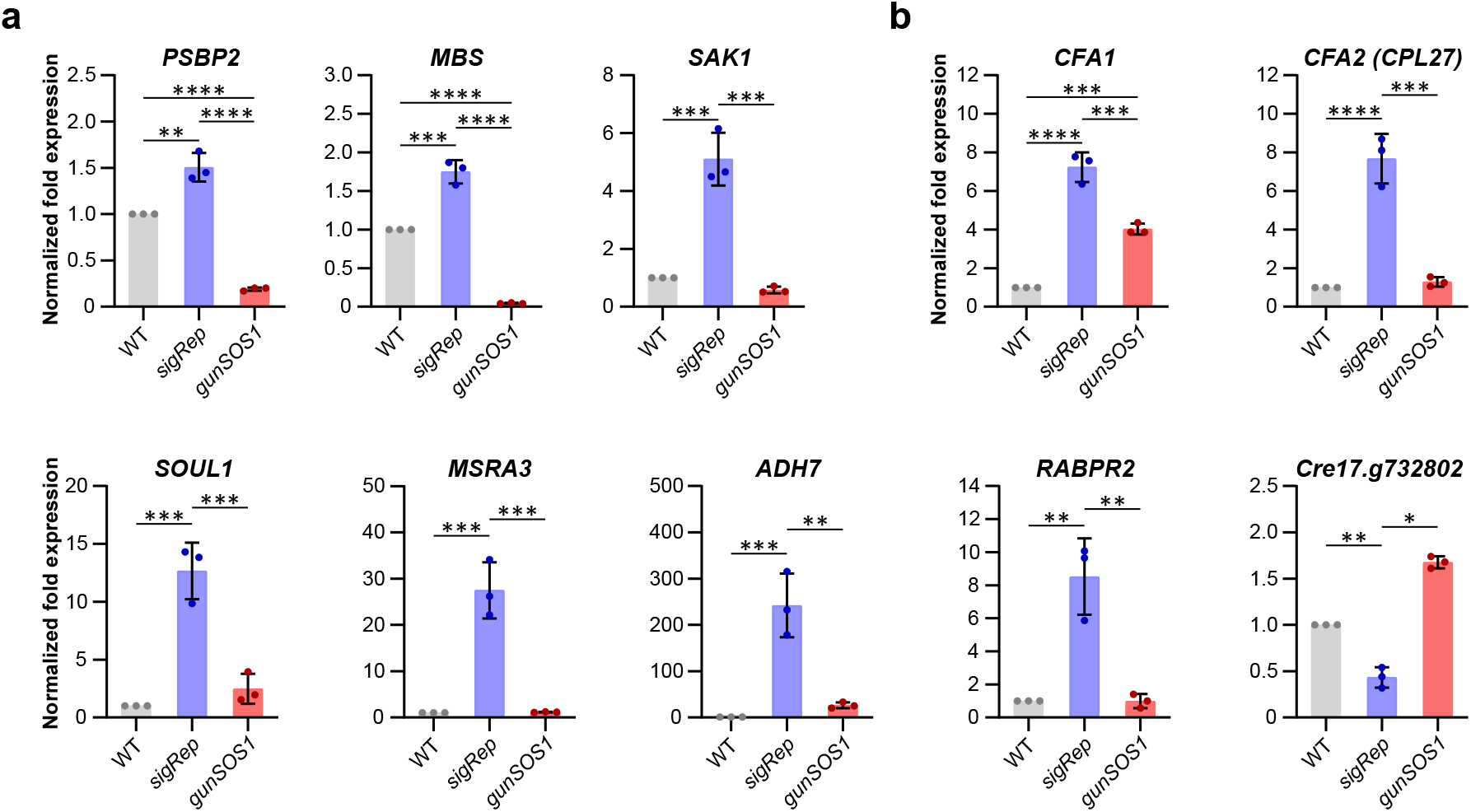
The qRT-PCR analysis of gene expression in *gunSOS1* compared to *sigRep*. **a** Expression of genes encoding proteinaceous components previously associated with the chloroplast retrograde signalling involving ^1^O_2_, *PSBP2*^15^, *MBS*^1^, and *SAK1*^6^. **b** Transcript levels of the selected genes showing attenuated expression in *gunSOS1* compared to *sigRep* in the context of the SAK1-transcriptome^16^, genes description in the text. Experiments were performed in biological replications (*n* = 3); results are presented as normalized fold expression (2^−ΔΔCt^, WT = 1); the error bars represent calculated ±SD. Significant differences were calculated using one-way ANOVA, pair-wise comparison with the Tukey’s post-hoc test (non-significant not shown), \**P* < 0.05, \*\**P* < 0.01, \*\*\**P* < 0.001, and \*\*\*\**P* < 0.0001.

### Altered metabolite content affects ^1^O_2_-signalling

We hypothesized that at least some of the metabolites showing increased or decreased content in *gunSOS1* compared to *sigRep*, attenuate or propagate ^1^O_2_-signalling, respectively. We selected fumarate and aconitate as representatives of either group. To test our hypothesis, metabolites were added separately to the liquid cultures at empirically determined sublethal concentrations, followed by incubation in the light to induce ^1^O_2_ generation by Proto. Subsequently, the *GPX5_cyt_* and *GPX5_cp_* transcript levels were used as readouts for the ^1^O_2_ signalling efficiency in *gunSOS1* and *sigRep*.

Exogenous application of fumarate significantly (*P* < 0.001, a detailed report from two-way ANOVA analyses is presented in Supplementary Tables 4 and 5) attenuated expression of both *GPX5_cyt_* and *GPX5_cp_* in a concentration-dependent manner in *sigRep* (Fig. 6a), which otherwise showed lower fumarate content compared to *gunSOS1* (Supplementary Fig. 5a) and had a functional ^1^O_2_-signaling (Figs. 1h,i and 6a). As expected, fumarate did not affect *GPX5* expression in *gunSOS1*. Application of 20 μM aconitate significantly (*P* < 0.0001) increased expression of *GPX5* in *gunSOS1* and even further in *sigRep* compared to untreated cells (*P* < 0.0001), but 100 μM had a quenching effect on the *GPX5* expression in *sigRep* (Fig. 6b, Supplementary Tables 6 and 7). This indicates that aconitate up to a certain threshold concentration promotes ^1^O_2_ signalling.

**Fig. 6.**
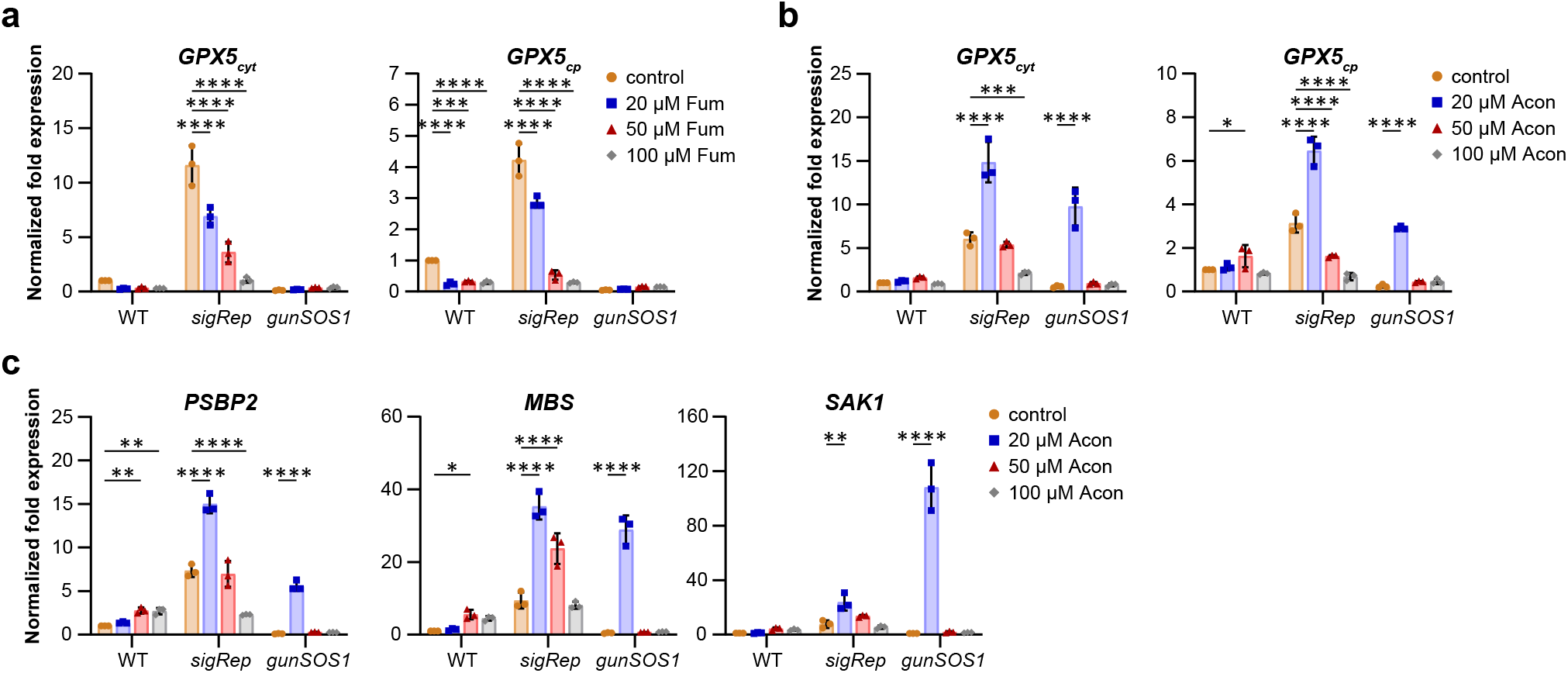
Effect of selected metabolites applied exogenously on ^1^O_2_-signalling inducing *GPX5_cyt_* or *GPX5_cp_* expression. **a** Fumarate (Fum) significantly (*P* < 0.0001) decreased *GPX5* expression in *sigRep* in concentration-dependent manner. Interaction between mutants and fumarate, as well as the mutant-dependent expression of *GPX5* were also significant, *P* < 0.0001 and *P* < 0.001, respectively. Fumarate had no effect on *GPX5*_cyt_ expression in *gunSOS1* or WT, but attenuating effect was observed for *GPX5_cp_* expression in WT (*P* ≤ 0.0001). Detailed reports from statistical analyses are presented in Supplemental Tables 4 and 5. **b** Aconitate (Acon) at 20 μM concentration rescued *GPX5* expression in *gunSOS1* and increased *GPX5* expression in *sigRep*. Statistical analysis showed that 20 μM aconitate significantly (*P* ≤ 0.0001) affected *GPX5_cyt_* expression in *gunSOS1* and *sigRep*, but 100 μM concentration decreased this expression in *sigRep (P* = 0.0001). An increase in *GPX5*_cp_ expression (*P* < 0.05) was also observed in WT upon treatment with 50 μM aconitate (Supplemental Tables 6 and 7). **c** Transcripts content for proteins necessary for the ^1^O_2_-signalling, *PSBP2, MBS*, and *SAK1* significantly (*P* < 0.01) increased upon treatment with 20 μM aconitate. An increase in *PSBP2* and *MBS* expression was also observed in WT treated with 50 μM aconitate. Detailed statistical analyses are presented in Supplemental Tables 8-10. The qRT-PCR experiments were performed in biological replications (*n* = 3); results are presented as normalized fold expression (2^−ΔΔCt^, untreated WT = 1); the error bars represent calculated ±SD. Significant differences were calculated using two-way ANOVA with Dunnett’s multiple comparison test and are indicated by asterisks (non-significant changes are not shown on the graph for clarity), \**P* < 0.05, \*\**P* < 0.01, \*\*\**P* < 0.001, and \*\*\*\**P* < 0.0001.

Subsequently, we determined the expression of *SAK1* upon feeding *gunSOS1* and *sigRep* with aconitate. Exogenous application of aconitate at 20 μM increased *SAK1* transcript abundance in *gunSOS1* more than 120 times compared to the untreated *gunSOS1* control, and exceeded values observed in *sigRep* subjected to the same treatment by a factor of 5 (Fig. 6c). Due to the increased *SAK1* expression in *gunSOS1* upon feeding with aconitate, we also determined the expression of *PSBP2* and *MBS* in the same conditions. The *PSBP2* response to aconitate was similar to *SAK1* in terms of the increased expression (Fig. 6c), but with less pronounced dependence on the metabolite concentration compared to *SAK1*. Although 20 μM aconitate increased *PSBP2* expression in *gunSOS1* by a factor of 48 compared to the untreated control, it was still 2.7 times lower compared to *sigRep* subjected to the same treatment (Fig. 6c). Similarly to *SAK1* and *PSBP2*, 20 μM aconitate also increased expression of *MBS* in *gunSOS1*, although values did not exceed those observed in *sigRep* subjected to the same treatment (Fig. 6c). Statistical analyses indicated that the effect of 20 μM aconitate on *SAK1, PSBP2* and *MBS* expression in *sigRep* and *gunSOS1*, interaction between mutants and aconitate, as well as the mutant-dependent expression of these genes were always significant (*P* < 0.001, Supplementary Tables 8-10).

### Specificity of the ^1^O_2_-signalling pathway(s) attenuated in *gunSOS1*

The profound effect of accumulating Tre6P on the transcriptome (Fig. 4a,b) and consequently the metabolism of *gunSOS1* (Fig. 3 and Supplementary Figs. 5 and 6) may have been indicative of a general photooxidative stress response being impaired in *gunSOS1*. To determine the specificity of this response, we compared *gunSOS1* and *sigRep* in terms of the expression of selected genes that are known to be induced by various reactive species or conditions causing photooxidative stress other than ^1^O_2_.

Upon H_2_O_2_ or organic *tert*-butyl hydroperoxide (*t*-BOOH) treatment, Blaby, et al. ^35^ observed an increase in the expression of the *MSD3* gene, encoding plastid-localised Mn SUPEROXIDE DISMUTASE 3. In our studies, we did not observe an increase in *MSD3* transcript in *sigRep* compared to WT (Supplementary Fig. 11a), which shows that *MSD3* expression is not inducible by ^1^O_2_ produced by Proto. However, based on qRT-PCR analysis, a 9-fold increase was observed in *gunSOS1* compared to *sigRep* (Supplementary Fig. 11a, see also RNA-seq in Supplementary Dataset 2). Upon H_2_O_2_ or *t*-BOOH treatment, Urzica, et al. ^36^ also observed induced expression of genes involved in the glutathione-ascorbate system, *GDP-L-GALACTOSE PHOSPHORYLASE* (*VTC2*) and *DEHYDROSASCORBATE REDUCTASE* (*DHAR1*). Based on our qRT-PCR results, *VTC2* transcript content was not changed in *sigRep* compared to WT, but an increase was observed in *gunSOS1* (Supplementary Fig. 11b and Supplementary Dataset 2). *DHAR1* expression was also stimulated in *gunSOS1* compared to *sigRep* (Supplementary Fig. 11c and Supplementary Dataset 2).

Similarly to *sak1* following treatment with rose bengal^16^, we observed increased expression of *GLUTATHIONE S-TRANSFERASE* (*GSTS1*) in *gunSOS1* compared to *sigRep* (Supplementary Fig. 11d and Supplementary Dataset 2). However, in another study increased expression of *GSTS1* was shown after treatment with acrolein, which suggests its transcriptional induction by reactive electrophile species (RES)^37^. Acrolein was also shown to induce *FSD1* encoding Fe SUPEROXIDE DISMUTASE (FeSOD)^37^, which was expressed both in *sigRep* and *gunSOS1* (Supplementary Dataset 1), and no significant difference could be observed in DEG analysis (Supplementary Dataset 2) or qRT-PCR (Supplementary Fig. 11e). It can be concluded that, despite impaired ^1^O_2_-signalling, *gunSOS1* retained the ability to respond to photooxidative stress caused by other ROS such as H_2_O_2_, organic peroxides, or RES.

## Discussion

ROS are formed as a by-product of biological redox reactions^38^ mostly in the mitochondria or chloroplasts^39,40^. Although excess ROS production can cause oxidative damage to cell components, ROS or the oxidation products play an important role in the signal transduction processes. Different retrograde signalling pathways have been proposed to involve also TBS-intermediates in plants and green algae (reviewed in^41,42^). While involvement of Mg-porphyrins in chloroplast retrograde signalling is now excluded^43,44^, in the present study we have demonstrated that ^1^O_2_ produced by the photosensitizing activity of Proto in the light triggers signalling cascades that alter nuclear gene expression in mutant that endogenously accumulate Proto.

The ^1^O_2_-induced signalling phenotype in the *gunSOS1* mutant was due to a lesion in the *TSPP1* gene. We demonstrated that TSPP1 is a functional phosphatase responsible for dephosphorylating Tre6P (Fig. 2a,d). *TSPP1* is the only representative of the classical plant *TPP* gene family in *C. reinhardtii*, contrasting with the large *TPP* gene families in angiosperms^45^. In addition, *C. reinhardtii* has one representative of the class I *TREHALOSE-6-PHOSPHATE SYNTHASE* (*TPS*) family (Cre16.g662350, hereafter *TSPS1*), and two members of the class II *TPS* family (Cre06.g278221, here *TSSP1* and Cre16.g686200, here *TSSP2*). Both class I and class II TPS proteins have glucosyltransferase and TPP-like domains, but only the class I TPS proteins have demonstrated TPS activity^46–48^. TSPP1 belongs to the haloacid dehalogenase superfamily of proteins and contains the characteristic DXDX(T/V) active site motif – _107_DYDGT_112_ – in which the initial Asp residue forms a phospho-acyl intermediate during catalysis^49^. TSSP1 also contains the complete active site motif (_592_DYDGT_597_), so we cannot exclude the possibility that the TSSP1 protein in *C. reinhardtii* has a phosphatase activity, which could explain the increased content of trehalose in *gunSOS1* compared to *sigRep* (Fig. 2c). Although the intracellular localizations of these proteins have not been determined experimentally in *C. reinhardtii*, PredAlgo analysis indicated possible localization of TSPP1 and TSSP1 in mitochondria, while TSPS1 and TSSP2 are likely to be cytosolic proteins, because they were not assigned to any specific organelle.

Correlation between *TSPP1* (Fig. 2b) and *GPX5* (Fig. 1b,c) expression may be indicating that low Tre6P content is necessary for efficient ^1^O_2_-signalling. However, the apparent negative effect of accumulating Tre6P or the absence of TSPP1 on ^1^O_2_-signalling is not direct and involves a complex metabolic reprogramming (Fig 4a,b) leading to altered metabolite content (Figs. 3 and Supplementary Figs. 5 and 6). Study on the contrasting phenotypes between *A. thaliana* overexpressing bacterial TPS, TPP, or TREHALOSE PHOSPHATE HYDROLASE (TPH) pointed to Tre6P, rather than trehalose, playing a signalling function^28^. Nevertheless, Tre6P was shown to be highly correlated with sucrose, leading to the proposal that it functions as a signal of sucrose status^26^. Tre6P was also shown to inhibit starch degradation in *A. thaliana* ^50,51^, which is also true in Tre6P-acumulating *gunSOS1* mutant of *C. reinhardtii* (Supplementary Fig. 6c). However, although our analyses revealed the sensitivity of ^1^O_2_-dependent retrograde signalling to metabolites, it is less clear if Tre6P or the TSPP1 protein itself, or both, play a direct role in metabolic reprogramming (Fig. 3 and Supplementary Figs. 5 and 6). TPP in *Z. mays* is encoded by *RAMOSA3* (*RA3*) and the *ra3* mutants showed reduced meristem determinacy^52^, without altering the Tre6P content compared to wild type plants^53^. Additionally, a catalytically inactive version of RA3 complemented the *ra3* phenotype, which revealed the “moonlighting” function of TPP, i.e. its function aside from the catalysis of Tre6P dephosphorylation^53^. Subsequent study also demonstrated “moonlighting” function of RA3 in carpel suppression^54^. Rather regulatory than a catalytic function was also proposed for TPP7 in *Oryza sativa*, due to its low activity *in vitro* and no apparent effect on Tre6P content in knockout mutants of TPP7^55^. In either case, although in our study the lack of functional TSPP1 clearly results in accumulation of Tre6P, we cannot exclude that the TSPP1 protein itself is involved in inducing changes to the nuclear gene expression (Fig. 4) and metabolic reprogramming (Fig. 3 and Supplementary Figs. 5 and 6).

Nonetheless, it was demonstrated that Tre6P acts as an inhibitor of SUCROSE-NON-FERMENTING (SNF)-RELATED PROTEIN KINASE1 (SnRK1) in developing tissues and that this is dependent on a so-far unidentified protein factor^56–58^. Tre6P also inhibits the activation of SnRK1 by SnRK1-activating kinases/geminivirus Rep interacting kinases^59^. SnRK1 belongs to the AMPK-SNF1-SnRK family of protein kinases, which is represented in all eukaryotes^60^. In plants, SnRK1 plays a central role in energy and metabolic homeostasis, and is activated during energy deficient conditions caused by stresses like nutrient starvation, pathogen attack, or ROS^61^. Baena-González, et al. ^62^ established approximately 1000 genes as markers of SnRK1 in *A. thaliana*, which indicates an extensive SnRK1-dependent transcriptional reprogramming. In *C. reinhardtii*, involvement of various SnRKs in responses to stress was observed during sulphur^63,64^ and nitrogen deprivation^65^, or cold stress^66^. Genome-wide analysis revealed the existence of 21 genes as potential orthologues of the plant SnRK α, β and γ/βγ subunits in *C. reinhardtii*. It was suggested that the proteins encoded by these genes play the same role in cell survival and stress response in *C. reinhardtii* as SnRKs in land plants^65^.

In analogy to *A. thaliana*, altered SnRKs activity was shown to cause metabolic remodelling also in algae ^66^. In our study, if the majority of changes to gene expression observed in *gunSOS1*, relative to *sigRep*, originate from accumulation of Tre6P and not the absence of TSPP1, based on previous studies, this leads to inhibition of one or more SnRKs (Fig. 7). Our analysis showed that altered gene expression in *gunSOS1* relative to *sigRep* (Supplementary Figs. 8-10) is directly responsible for altered fumarate and aconitate content in *gunSOS1* compared to *sigRep*. Thus, Tre6P accumulating in *gunSOS1* is not directly involved in ^1^O_2_-signalling but controls other processes in the cell which affect ^1^O_2_-signalling more directly (Fig. 7).

**Fig. 7.**
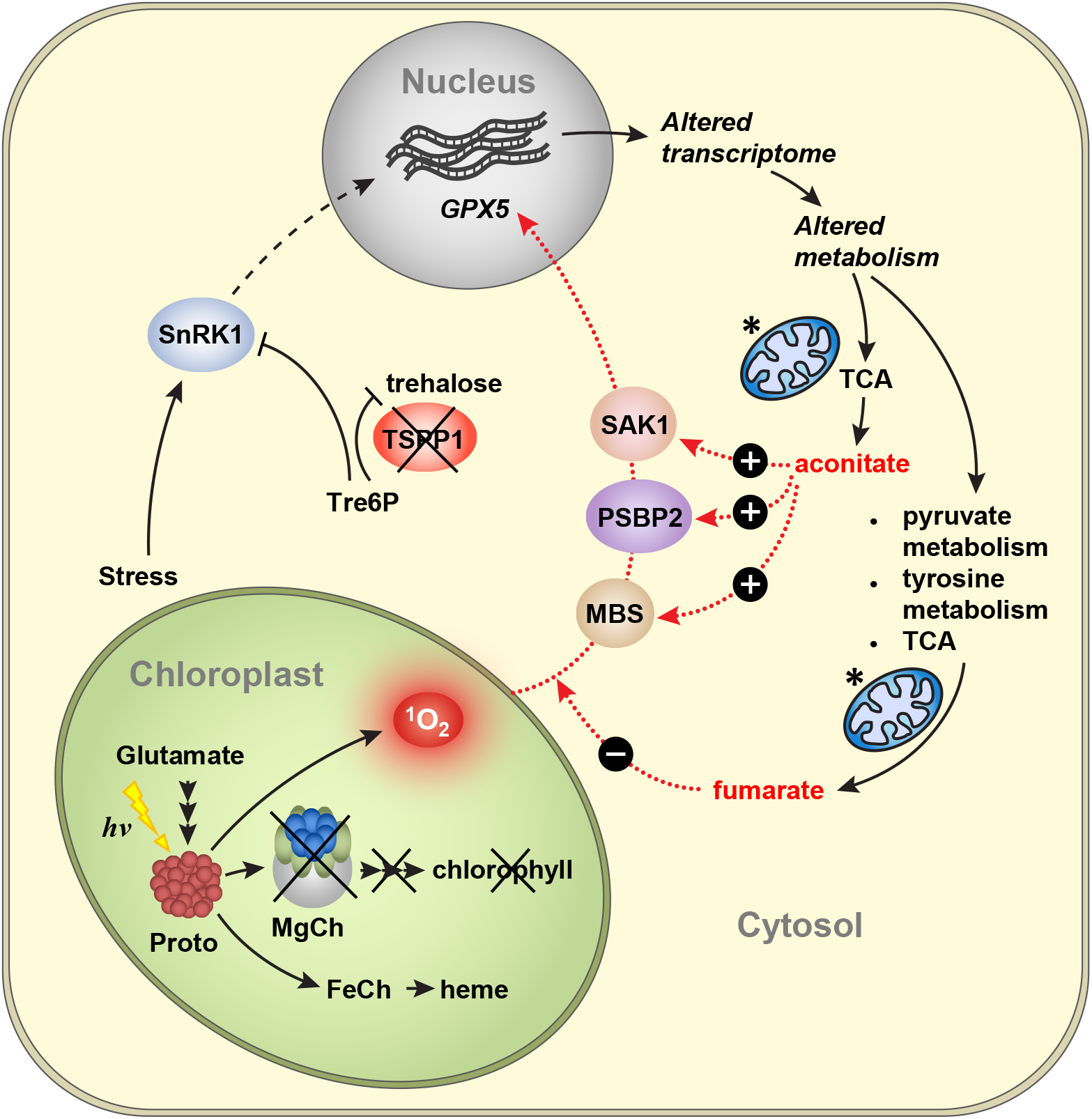
Model of the chloroplast ^1^O_2_-induced retrograde signalling depending on the metabolites content. Lack of a functional Mg-chelatase (MgCh) leads to accumulation of protoporphyrin IX (Proto), which in light generates ^1^O_2_. Absence of TSPP1 results in accumulation of trehalose 6-phosphate (Tre6P). In plants, Tre6P has an inhibitory effect on SnRK1^56–58^, while altered SnRKs activity is also involved in stress responses in *C. reinhardtii*^63–66^. Blocked SnRK1 signalling pathway leads to the inadequate transcriptomic response to stress and disturbed metabolites content. Fumarate accumulated in *gunSOS1* relative to *sigRep* due to the altered expression of genes involved in pyruvate metabolism, tyrosine metabolism and the TCA cycle. Exogenous application of fumarate had an inhibitory effect on ^1^O_2_-signaling in *sigRep*, while no change was observed in *gunSOS1* and WT, relative to their respective non-treated controls. Accumulation of fumarate and disturbed TCA cycle may lead to deficiency in aconitate. The proteinaceous components, PSBP2, MBS, and SAK1, previously demonstrated to be required for ^1^O_2_-signalling are shown. Exogenous application of aconitate promoted expression of *PSBP2, MBS*, and *SAK1*, and consequently also *GPX5*, which ultimately indicated the rescue of ^1^O_2_-signaling in *gunSOS1*. Intracellular localisation of TSPP1, SnRK1, and PSBP2 depicted in the Figure has not been determined experimentally and has only an illustrative character. Mitochondria are marked by an asterisk.

We demonstrated that accumulation of fumarate (Fig. 3 and Supplementary Fig. 5a) is responsible for attenuating ^1^O_2_-signalling (Fig. 6a, see also model in Fig. 7). Fumarate is a well-recognized oncometabolite in mammalian cells^67^, associated with development of tumours by competitive inhibition of 2-oxoglutarate-dependent oxygenases, including HYPOXIA INDUCIBLE FACTOR (HIF) hydroxylases, leading to stabilisation of HIF and activation of oncogenic HIF-dependent pathways^68^. In fact, accumulation of fumarate in human cells was linked to an aggressive variant of hereditary kidney cancer^69^. In mice, fumarate was also shown to directly modify some proteins by succination of cysteine residues to form 2-succinocysteine derivatives^70^. Succination of three cysteines crucial for iron-sulphur cluster binding was identified in mitochondrial ACONITASE2 (ACO2) in a *fumarate hydratase 1* knockout (Fh1KO) mouse embryonic fibroblast (MEF) cell line^70^. Analysis of tryptic peptides derived from ACO2 in Fh1KO MEFs, indicated succination of Cys385. Another tryptic peptide of ACO2 was identified as a mixture of two isomers in Fh1KO MEFs, which showed succination at Cys451 or Cys448^70^. *In vitro* experiments indicated that succination of these cysteine residues in ACO2 leads to its inhibition^70^. Equivalent cysteine residues are conserved in *C. reinhardtii* ACH1, which in part explains the aconitate deficiency in *gunSOS1* (Fig. 3 and Supplementary Fig. 6a). Thus, the diminished expression of *ACH1* in *gunSOS1* compared to *sigRep* (Supplementary Fig. 10a,b) and plausible inactivation of ACH1 by fumarate, together may constitute factors leading to aconitate depletion in *gunSOS1*.

The rescued expression of the genes encoding PSBP2^15^, MBS^17^, and SAK1^16^ (Fig. 6c), consequently rescued ^1^O_2_-signaling and the expression of *GPX5* (Fig. 6b) upon exogenously applied aconitate is intriguing, but the underlying mechanism remains unknown. There is no information in the literature regarding involvement of aconitate in the signaling. However, substantial amount of data indicated the key role of aconitase (EC 4.2.1.3) in responses triggered by ROS, due to the presence of the iron-sulfur cluster^71,72^. In animal cells, aconitase redox-regulated moonlighting function modulates biosynthesis of proteins carrying Iron Response Element (IRP) in their mRNA^72–75^. The dual function of aconitase is possible due to the reversible redox-dependent post-translational modifications (reviewed in^74^). Three cysteines crucial for binding the iron-sulphur cluster in mitochondrial ACONITASE2 (ACO2) in mouse, Cys385, Cys448, and Cys451^70^, are also present in ACH1 of *C. reinhardtii*, Cys426, Cys489, and Cys492, respectively.

Aconitase emerges as a factor involved in stress response also in plants. In *A. thaliana*, ACONITASE (ACO) is found in three isoforms, while ACH1 is the only enzyme annotated at the JGI portal as aconitase in *C. reinhardtii*. ACO3 in *A. thaliana* serves both as a target and mediator of mitochondrial dysfunction signaling, and it was shown to be critical for stress response in leaves^76^. Phosphorylation of ACO3-Ser91 contributes to the UV-B and mitochondrial complex III inhibitor antimycin A-induced stress tolerance. ACH1 of *C. reinhardtii* contains four serine residues within a widely conserved eukaryotic phosphorylation motif R-x-x-S^77^, Ser118, Ser211, Ser284, and Ser439, so their phosphorylation is likely. ACO3 was demonstrated to be part of the mitochondrial dysfunction response, which is dependent on the signalling involving transcription factor NAC DOMAIN CONTAINING PROTEIN 17 (ANAC017). ANAC017 is considered as a master regulator of the retrograde signalling and cellular stress response with mitochondria acting as central sensors, but also during repression of the chloroplast function^78^. Based on the genome-wide association study concerning single nucleotide polymorphism, it was determined that promoter of ACO3 binds ANAC017^79^.

ACO3 plays an important role in acclimation to submergence^79^ and it was proposed to regulate the stability of CHLOROPLAST SUPEROXIDE DISMUTASE2 (SOD2) mRNA^80^. Both submergence-related hypoxia and reoxygenation during desubmergence are accompanied by increased ROS generation and oxidative stress^81,82^. The *aco3* knockout mutant and *ACO3* overexpressing lines (*ACO3OE*) showed altered stress signalling correlated with decreased and increased stress tolerance, respectively. Furthermore, a decreased expression of FUMARASE 2 (FUM2) was noted in *ACO3OE* lines in the control conditions, and it decreased in *aco3*, *ACO3OE*, and in WT after submergence and desubmergence^79^. The JGI portal blast-search analysis of *A. thaliana* FUM2 against *C. reinhardtii* proteome indicated, that the protein showing the highest similarity in amino acid sequence is the FUM2, which showed increased expression in *gunSOS1* compared to *sigRep* (Supplementary Fig. 8c, see also Supplementary Datasets 2 and 3). These findings suggest a reciprocal interplay between fumarate and aconitate in response to stress.

The decreased expression of *ACH1* does not affect the content of citrate and isocitrate, but only deficiency in aconitate was observed in *gunSOS1* compared to *sigRep*. Based on our results and current knowledge, it can be hypothesized that not aconitate, but ACH1 is the crucial player in ^1^O_2_-signaling in *C. reinhardtii*, while accumulation of fumarate deactivates ACH1, thereby impairing oxidative stress sensing and ^1^O_2_-signalling. Chloroplasts and mitochondria are biochemically connected^83,84^, while the function of the components localized in either compartment has been shown to be integrated into the functional signaling^85–89^. Therefore, it seems plausible that the mitochondrial function participates, or is even required for functional chloroplast-emitted ^1^O_2_-signaling. The chloroplast ^1^O_2_-signal might be providing an input into the overall signaling network, which is subsequently integrated or dependent on the mitochondrial retrograde signaling and other internal or external stimuli.

Our results regarding increased content of fumarate and decreased aconitate in *gunSOS1* compared to *sigRep* (Fig. 3 and Supplementary Figs. 5 and 6) as well as feeding experiments (Fig. 6) indicate, that there is a metabolic signature conditioning ^1^O_2_-signalling. However, it should be noted that the extent to which exogenously applied fumarate and aconitate are taken up into the cells was not determined. Nevertheless, treatment with fumarate and aconitate, both have a very clear impact on gene expression (Fig. 6), which would not take place, if these metabolites were not taken up or sensed by the cells. Although, there are no known receptors detecting the presence of these metabolites outside of the cell, their existence cannot be excluded at this point. In either case, it was shown that altered chloroplast redox conditions result in changes in the metabolome in *A. thaliana*, reflected also by the reallocation of energy resources^90^. Importantly, although our study shows that the metabolic configuration of the cell is essential for ^1^O_2_-signalling, this does not exclude the involvement of protein components in this signalling network (Fig. 7). On the contrary, our findings support the direct involvement of proteins in ^1^O_2_-signalling, but we demonstrated that expression of *PSBP2*^15^, *MBS*^17^, and *SAK1*^16^ depends on the metabolic status (Fig. 5a and Fig. 6c). The overlap between the *gunSOS1* and the SAK1-dependent transcriptomes (Fig. 5b) as well as other defined ROS or RES-dependent signalling pathways being functional (Supplementary Fig. 11) indicate specificity of the impaired ^1^O_2_-signalling in *gunSOS1*. Our results indicate that fumarate and aconitate affect the expression of *PSBP, MBS*, or *SAK1*, which subsequently convey information about the metabolic status of the cell to the nucleus and trigger specific responses (Fig. 7). It is not clear yet how the specificity of the signalling is achieved, because the mechanisms by which metabolites affect the expression of these proteins, as well as downstream components of these signalling pathways, remain unknown. Additionally, the protein components involved in ^1^O_2_-signalling may be subject to additional regulatory mechanisms, such as phosphorylation, as proposed for SAK1 by Wakao, et al. ^16^.

We have shown that retrograde signalling triggered by ^1^O_2_, which is mostly generated in the chloroplast, strictly depends on the mitochondrial metabolic status. PROLYL-tRNA SYNTHETASE (PRORS1) in *A. thaliana*, a component of the organellar gene expression machinery indicated in the previous study that mitochondria may contribute to chloroplast retrograde signalling^91^. PRORS1 is targeted both to the plastid and to mitochondria, but downregulation of specific photosynthesis-associated nuclear genes was observed only in a plastidial *prpl11* and mitochondrial *mrpl11* double mutant, but not in *prpl11* or *mrpl11* single mutants^91^. Another study showed that the ROS-dependent signals from chloroplast and mitochondria are integrated at the RADICAL-INDUCED CELL DEATH 1 (RCD1) protein located in the nucleus^89^. RCD1 suppresses transcription factors ANAC013 and ANAC017, which mediate the ROS-signal from mitochondria, while chloroplast regulation of RCD1 takes place through 3’-phosphoadenosine 5’-phosphate. It was proposed that RCD1 may function at the intersection of mitochondrial and chloroplastic retrograde signalling pathways^89^.

The retrograde signalling pathways identified so far assume the existence of separate signalling routes. However, chloroplast retrograde signalling involves a network of different but inter-connected mechanisms, and observation of specific signalling pathways within the network depends on the experimental conditions. Pharmacological or genetic interventions that disrupt specific pathways can be used to characterize alternative pathways. In the manuscript presented here, we report that the metabolic status of the cell, reflecting levels of Tre6P, and components of the mitochondrial TCA cycle, are integral factors in the ^1^O_2_-dependent retrograde signalling network in *C. reinhardtii*. It remains a matter of debate whether Tre6P or TSPP1 protein itself plays a direct role in the signalling and metabolic reprogramming. Additionally, further study should elucidate whether aconitate or ACH1 is the key player in the ^1^O_2_-signaling. Nevertheless, our results shed an important light on the signalling processes triggered by ^1^O_2_, by revealing their sensitivity to metabolites and thereby their potential modulation by cytosolic and mitochondrial metabolism.

## Methods

### *Chlamydomonas reinhardtii* cultures and genetic manipulations

All strains were cultivated heterotrophically in Tris-acetate-phosphate (TAP) medium in the dark. Additionally, starch measurements were performed in TAP, Tris-phosphate (TP) medium without acetate, or nitrogen-depleted (TAP-N) medium. Experiments were performed in TAP medium in the dark or upon a shift to 20 μmol photons m^−2^ s^−1^ light. Sampling of three separate cultures grown in parallel in the same conditions was considered as biological triplicates. Three strains used in this study were described elsewhere, *chlD-1* in von Gromoff, et al. ^18^, *chlD-1/GUN4* in Brzezowski, et al. ^19^, and wild type (4A+) in Dent, et al. ^92^.

To generate the *GPX5-ARS2* construct (Supplementary Fig. 2), the *GPX5* regulatory region (*GPX5* 5’ RR) was amplified by PCR using forward and reverse primers carrying XhoI and EcoRV restrictions sites, respectively (Supplementary Table 11). The obtained fragment was subcloned and ligated between the XhoI/EcoRV sites of the pSL18 vector^93^, replacing the existing promoter region of the gene encoding the PSAD and positioning *GPX5* 5’RR in reverse orientation with respect to the paromomycin resistance cassette (paroR^94^. The DNA fragment carrying paroR and *GPX5* 5’ RR was excised by restriction digestion with KpnI and EcoRV and ligated into the corresponding restriction sites of pJD54^95^, which carries a copy of the promoterless version of the gene encoding ARYLSULFATASE 2 (ARS2), which allowed *ARS2* expression to be controlled by the *GPX5* 5’RR (Supplementary Fig. 2). The *GPX5-ARS2* construct was verified by sequencing.

The *sigRep* strain was produced in the *chlD-1/GUN4* background by transformation with a *GPX5-ARS2* construct. Transformant selection was performed on TAP agar plates with 10 μg mL^−1^ paromomycin in dark, followed by a screen for *GPX5-ARS2* inducibility by ^1^O_2_. The *sigRep* strain showed low ARS2 activity in the dark and high activity in the light (Fig. 1A) and was selected for further applications. The *gunSOS1* mutant was generated by random insertional mutagenesis performed on *sigRep* using a bleomycin resistance cassette (ble^R^) isolated from the pMS188 vector^96^ using NotI and KpnI. Following mutagenesis, selection was performed on TAP agar plates containing 15 μg mL^−1^ zeocin in the dark. More than 800 obtained colonies were screened for decreased or not detectable ARS2 activity in the light and nine mutants that showed impaired inducibility of *GPX5-ARS* expression in light were selected for further analysis. Restriction enzyme site-directed amplification PCR^97^ allowed us to identify the genomic DNA flanking the ble^R^ insertion sites in five out of the nine mutants. Only analysis of the *gunSOS1* mutant is presented here. Rescue of *gunSOS1* (*tspp1*) was conducted with 5242 bp fragment of genomic DNA carrying 3241 bp *TSPP1* amplified by PCR (see Supplementary Table 11 for primers) using bacterial artificial chromosome PTQ5987 (Clemson University Genomics Institute, Clemson, SC, USA) as a template. The amplified DNA fragment included 1500 bp upstream and 501 bp downstream of the annotated *TSPP1*. Wild-type *TSPP1* was introduced into *gunSOS1* by co-transformation with a spectinomycin resistance cassette isolated from the pALM32 vector^98^ with AleI and KpnI endonucleases. Transformant selection was performed on TAP agar plates supplemented with 100 μg mL^−1^ spectinomycin in the dark. All genetic transformations were performed by electroporation.

### Arylsulfatase activity assay

To assess the level of *GPX5-ARS2* expression in transformed cells, enzymatic assays for arylsulfatase activity^23^ were performed essentially as described before^15^. Cells were spotted onto agar-solidified TAP medium plates and cultured in either light or dark conditions. Arylsulfatase is expected to be secreted into the medium if *GPX5-ARS2* is expressed. After removal of the cells, plates were flooded with detection solution containing 0.1 mg mL^−1^ 2-naphthyl sulfate (potassium salt; Santa Cruz Biotechnology, Inc., Dallas, TX, USA) as a chromogenic substrate coupled with 1 mg mL^−1^ tetrazotized-o-dianisidine chloride (Fast Blue B salt, Santa Cruz Biotechnology, Inc., Dallas, TX, USA). Following 1 h incubation, purple spots appearing on the agar plates identified expressed ARS2.

### Analysis of protoporphyrin IX content

The Proto content was analysed by High Pressure Liquid Chromatography (HPLC) essentially as described in Czarnecki, et al. ^99^, with modified sample preparation for *C. reinhardtii*^19^. In short, cultures were grown in the dark and transferred to 20 μmol photons m^−2^ s^−1^ light for 2 h. Samples containing 1.2 × 10^8^ cells were centrifuged at 3000 x *g* for 5 min at 4°C and the pellets were snap-frozen in liquid N_2_. Proto was extracted in cold (−20°C) acetone/0.1 M NH_4_OH (9/1, v/v) in a three-step cycle of resuspension and centrifugation. Proto analysis was performed using a Nova-Pak C18 column (Waters, 3.9 × 150 mm, 4 μm, at 20 °C). The results were normalized to pmol/10^6^ cells.

### RNA isolation, qRT-PCR, and RNA-seq

The total RNA was isolated, after a shift from dark to 20 μmol photons m^−2^ s^−1^ light for 2 h, using TRIzol Reagent (Thermo Fisher Scientific, Waltham, MA, USA), according to the manufacturer’s protocol. RNA quality was assessed by electrophoresis on a 1% (w/v) agarose gel, while quantity was determined using a Nanodrop 2000 (Thermo Fischer Scientific, Waltham, MA). Aliquots of 2 μg RNA were treated with DNase and RiboLock RNase inhibitor (Thermo Fisher Scientific, Waltham, MA, USA) and used to synthesize cDNA with RevertAid Reverse Transcriptase (Thermo Fisher Scientific, Waltham, MA, USA) and oligo(dT)_18_ primer. Transcript analysis by qRT-PCR were performed using 2× ChamQ Universal SYBR qPCR Master Mix (Viazyme Biotech Co., Ltd., Nanjing, China) and a CFX96-C1000 96-well plate thermocycler (Bio-Rad, Hercules, CA, USA), the 18S rRNA was used as a reference gene^25^. All primers were designed using Primer3Plus (http://primer3plus.com), and are listed in Supplementary Table 11.

RNA-sequencing was performed on total RNA samples isolated from biological triplicates of *sigRep* and *gunSOS1* after the shift from dark to 20 μmol photons m^−2^ s^−1^ light for 2 h. Library preparation, sequencing, and analysis services were commercially provided by Novogene Europe (Novogene (UK) Company Ltd., Cambridge, UK) and performed using an Illumina NovaSeq 6000 platform operated in 150 bp pair end sequencing mode, with a sequencing depth of 20 million reads per sample. Six gigabases of sequencing data per library was filtered for high-quality reads, which were mapped to the *C. reinhardtii* v5.5 (Department of Energy JGI). Estimated expression was obtained in fragments per kilobase of transcript sequence per millions base pairs sequenced (FPKM). Biological replicates were averaged to obtain sample expression estimate. Differential expression analysis between two mutants (three biological replicates per mutant) was performed using DESeq2.v1.20.0, which provides statistical routines for determining differential expression using a model based on the negative binomial distribution. The resulting *p* values were adjusted using the Benjamini and Hochberg’s approach for controlling the False Discovery Rate (FDR). Genes with a log_2_FoldChange ≥ 2 and adjusted *p* value (padj) < 0.05 found by DESeq2.v1.20.0 were defined as differentially expressed (DEGs). Gene Ontology (GO) analysis and Kyoto Encyclopedia of Genes and Genomes (KEGG; www.genome.jp/kegg/pathway.html) pathway analysis were conducted to identify DEGs at the biologically functional level using clusterProfiler R package. GO terms and KEGG pathways with a padj < 0.05 were considered to be significantly enriched.

### Metabolite analysis

Cultures were grown in TAP in the dark until they reached mid-log phase of 3 × 10^6^ cells mL^−1^. Samples were always normalized to contain 6 × 10^7^ cells, all centrifugation steps were carried at 3000 × *g* at 4°C for 5 min, followed by snap-freezing in liquid N_2_. Samples from dark conditions were collected immediately, while the remainder of each culture was transferred to 20 μmol photons m^−2^ s^−1^ light. Subsequent sampling took place at 15-min intervals, up to 1 h. Metabolites were extracted using chloroform-methanol as described by Lunn et al. (2006). Tre6P, other phosphorylated intermediates and organic acids were measured by anion-exchange HPLC coupled to tandem mass spectrometry (LC-MS/MS) as described in Lunn et al. (2006) with modifications as described in Figueroa, et al. ^100^. Sugars and sugar alcohols were measured by LC-MS/MS as described in Fichtner, et al. ^101^. Obtained results were normalized to pmol/10^6^ cells.

For starch analysis, cultures were grown in TAP in the dark until they reached mid-log phase of 3 × 10^6^ cells mL^−1^. For starch determination cultures were either transferred in TAP to the light (20 μmol photons m^−2^ s^−1^) for 2 h before harvesting, or centrifuged cells were resuspended in Tris-phosphate (TP) medium (without acetate), keeping the same cell concentration, and transferred back to the dark for 24 h. Subsequently, cultures were transferred to 20 μmol photons m^−2^ s^−1^ for 2 h before harvesting. A similar protocol was applied for starch analysis in TAP devoid of N (TAP-N), except that cells were cultivated in TAP-N for 3 days in the dark, followed by exposure to 20 μmol photons m^−2^ s^−1^ for 2 h. For each sampling, 5 × 10 cells were centrifuged and lyophilized. Starch content was determined by enzymatic degradation and glucose quantification following the protocol of DuBois, et al.^102^.

### Protein extraction and immunoblot analysis

Cultures were grown in the dark, followed by a transfer to 20 μmol photons m^−2^ s^−1^ for 3 h. Cells were pelleted by centrifugation and total proteins were extracted in 400 μL buffer containing: 56 mM Na_2_CO_3_, 56 mM DTT, 2% (w/v) SDS, 12% (w/v) sucrose and 2 mM EDTA, pH 8.0. Proteins were quantified using Pierce™ BCA Protein Assay Kit (Thermo Fisher Scientific, Waltham, MA, USA) and separated by SDS-PAGE on a 12% polyacrylamide gel, followed by transfer by electroblotting to nitrocellulose membrane (GE Healthcare, Chicago, IL, USA). TSPP1 was detected using a rabbit crude antiserum (dilution 1:500) raised by inoculation with a TSPP1-specific peptide (VEWSKSDSNGWRAKPC) against *C. reinhardtii* TSPP1 (calculated MW 42 kDa). GPX5 was detected with a commercially available antibody (AS15 2882, dilution 1:1000) obtained from Agrisera (Vännäs, Sweden). CHLI1 antibody (PHY5510S, dilution 1:1000) was purchased from PhytoAB (San Jose, CA, USA). For application, all antibodies were diluted in CrossDown buffer (AppliChem, AppliChem GmbH, Darmstadt, Germany). The secondary antibody (AS09 602, goat anti-rabbit IgG, dilution 1:10,000) conjugated to horseradish peroxidase was obtained from Agrisera. The immunoblotting signals were detected using a CCD camera (Intas Biopharmaceuticals, Ahmedabad, India) after application of enhanced chemiluminescence detection kit (Clarity™ Western ECL Substrate; Bio-Rad, Hercules, CA, USA.

### Chemical treatments of *C. reinhardtii* cells

Fumarate (sodium fumarate dibasic) and aconitate (*cis*-aconitic acid) were obtained from Sigma-Aldrich (Sigma-Aldrich Chemie GmbH, Taufkirchen, Germany), dissolved in H_2_O to a stock solution of 100 mM and added individually to the mid-log phase (3 × 10^6^ cells mL^−1^) cultures to final concentrations of 20, 50, and 100 μM, followed by exposure to light for 2 h.

## Supporting information

Al Youssef et al_Supplementary Figures

Al Youssef et al_Supplementary Tables

Al Youssef et al_Supplementary Dataset 1

Al Youssef et al_Supplementary Dataset 2

Al Youssef et al_Supplementary Dataset 3

## Acknowledgements

This work was supported by the Deutsche Forschungsgemeinschaft (DFG-GR 936/20-2 to B.G. and P.B.), by the Agence Nationale de la Recherche (ChloroPaths: ANR-14-CE05-0041-01 to X.J.), and by the Max Planck Society (R.F. and J.E.L.).

## Author Contributions

P.B. designed the research. P.B. and W.A.Y. performed most of the experiments. R.F. and J.E.L. performed the LC-MS/MS measurements. M.S.-S. and X.J. performed the starch quantification. P.B., B.G., and J.E.L. analysed the data. P.B. wrote the manuscript. P.B., B.G., J.E.L., and X.J. revised the manuscript. All authors discussed the results and commented upon the manuscript.

## Data availability

All data relevant for interpretation of this study are presented in the article and supplementary material. Any further information is available from the corresponding author upon reasonable request.

