## Supplementary material for "Singlet oxygen-induced signalling depends on the metabolic status of the Chlamydomonas cell": Al Youssef et al_Supplementary Figures

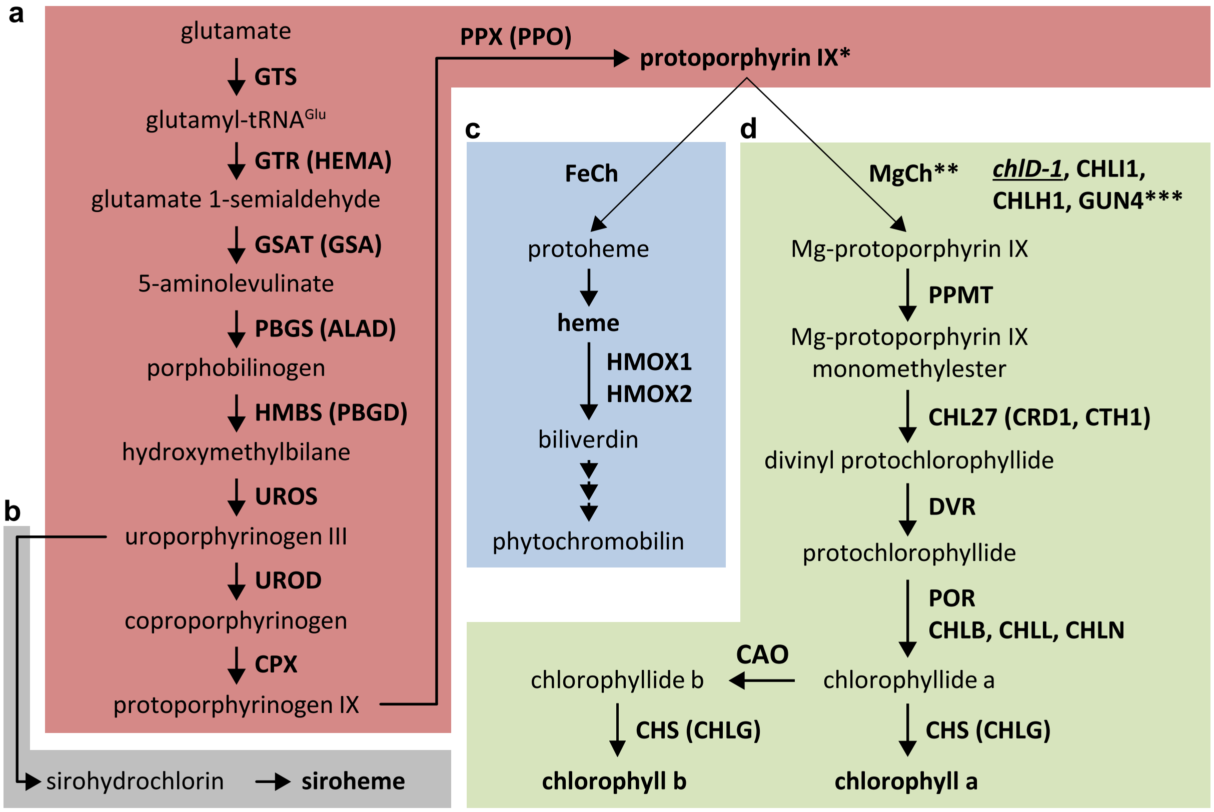


**Supplementary Fig. 1 Schematic representation of tetrapyrrole biosynthesis pathway.** **a** Common steps from glutamate to protoporphyrin IX (Proto); Proto is marked by an asterisk. **b** The siroheme biosynthesis branch. **c** Heme biosynthesis branch, from ferrochelatase (FeCh) to heme. Subsequent steps of heme catabolism from biliverdin to formation of phytochromobilin are not shown in detail. **d** Steps from Mg-chelatase (MgCh) to chlorophyll formation. MgCh responsible for inserting $Mg^{2+}$ into Proto is marked by a double asterisk. GUN4 protein involved in MgCh function is marked by a triple asterisk. Mutation in one of the subunits of MgCh, *chlD-1* is underlined.

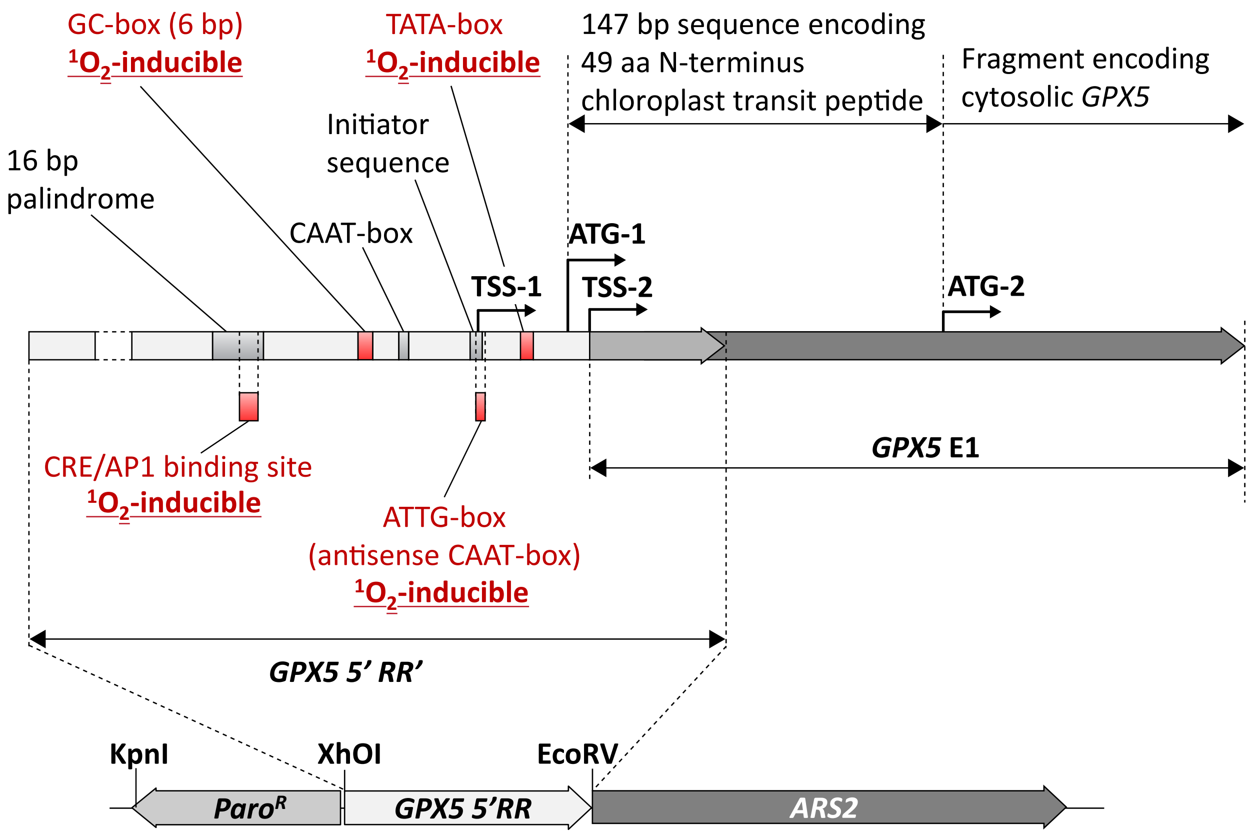


**Supplementary Fig. 2 Schematic representation of the *GPX5-ARS2* reporter gene construct.** Upper panel, the *GPX5* gene encodes two proteins, one targeted to the chloroplast equipped with the chloroplast transit peptide (*GPX5*$_{cp}$), and the cytosolic one (*GPX5*$_{cyt}$). Alternative transcription (TSS) and translation start sites (ATG) are indicated. The $^1O_2$-inducible *cis*-elements in *GPX5* regulatory region (*GPX5'RR*) are indicated, based on Fischer et al.[25]. Lower panel, fusion of *GPX5'RR* with *ARS2*, and the paromomycin resistance cassette (Paro$^R$) in reverse orientation is indicated.

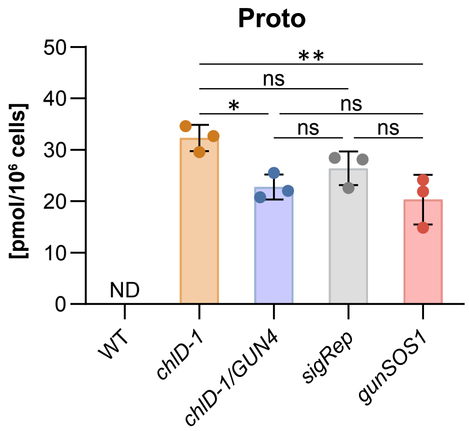


**Supplementary Fig. 3 Analysis of Proto content in *gunSOS1*.** Mutant impaired in $^1O_2$-signaling showed similar Proto accumulation in light compared to sigRep; ND, not detectable. Experiments were performed in biological replications ($n$ = 3); the error bars represent calculated ±SD. Significant differences were calculated using one-way ANOVA, pair-wise comparison with the Tukey's post-hoc test, non-significant (ns), *$P < 0.05$ and **$P < 0.01$. All mutants unable to synthesize chlorophyll showed significant accumulation of Proto compared to WT ($P < 0.0001$, not shown).

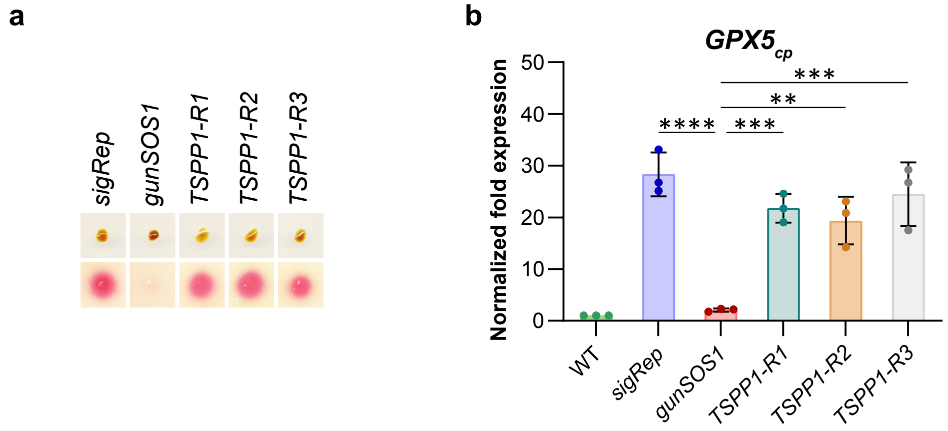


**Supplementary Fig. 4 Rescue of the $^1O_2$-signalling in *gunSOS1* by introduction of the wild-type *TSPP1*. a** Arylsulfatase assay showed higher activity of ARS2 in rescued strains, *TSPP1-R1*, *-R2*, and *-R3* compared to *gunSOS1*. **b** Expression of GPX5$_{cp}$ in the rescued strain compared to *gunSOS1* and *sigRep*. Results are presented as normalized fold expression ($2^{-\Delta\Delta Ct}$, WT = 1); experiments were performed in biological replications ($n = 3$); the error bars represent calculated ±SD. Significant differences were calculated using one-way ANOVA, pair-wise comparison with the Tukey's post-hoc test (non-significant not shown), $^{**}P < 0.01$, $^{***}P < 0.001$, and $^{****}P < 0.0001$. Statistical comparison between mutants and WT are not shown for clarity.

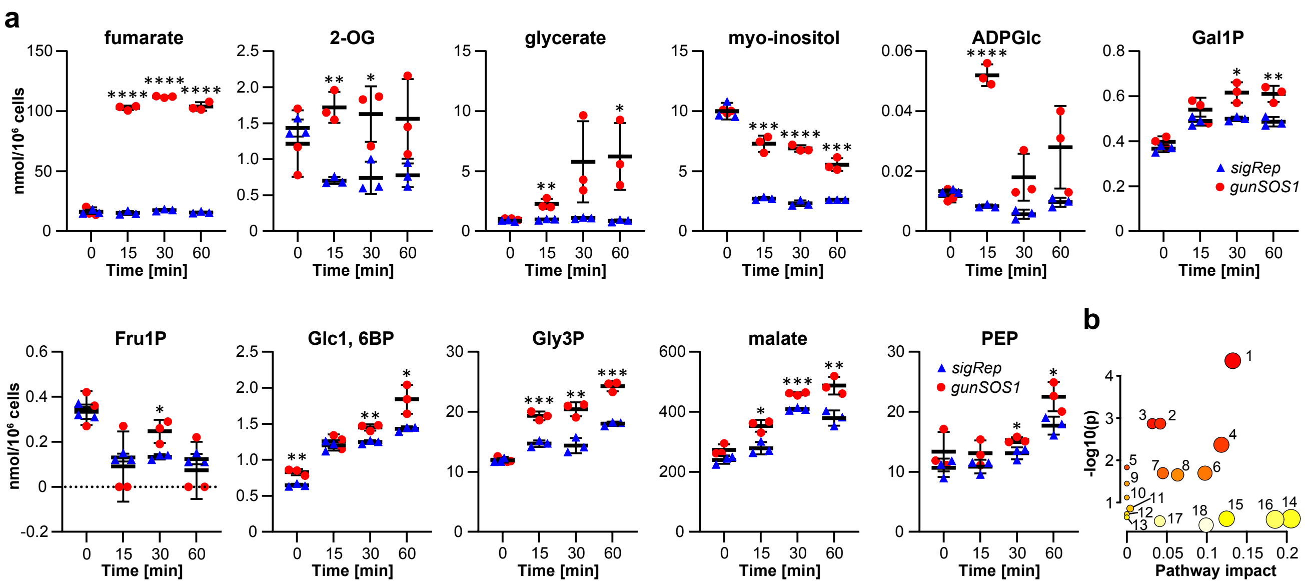


**Supplementary Fig. 5 Metabolites with significantly increased content in *gunSOS1* compared to *sigRep* in the light.** **a** LC-MS/MS analysis; fumaric acid (fumarate), oxoglutaric acid (2-OG), myo-inositol, ADP-glucose (ADPGlc), galactose 1-phosphate (Gal1P), fructose 1-phosphate (Fru1P), alpha-D-glucose 1,6-bisphosphate (Glc1, 6BP), glycerol 3-phosphate (Gly3P), phosphoenolpyruvic acid (PEP), and L-malic acid (malate). Measurements were performed in biological triplicates ($n$ = 3), horizontal bars represent the calculated mean, vertical error bars represent calculated ±SD; significant differences were calculated using two-tailed Student's $t$-test and are indicated by asterisks (non-significant not shown), *$P < 0.05$, **$P < 0.01$, ***$P < 0.001$, and ****$P < 0.0001$. **b** Impact of accumulating metabolites on cell metabolism analysed using MetaboAnalyst 5.0 (https://www.metaboanalyst.ca), sorted by the $P$ value; 1. tricarboxylic acid cycle (TCA cycle; $P$ = 4.41E-05), 2. pyruvate metabolism ($P$ = 0.00136), 3. starch and sucrose metabolism ($P$ = 0.00136), 4. glyoxylate and dicarboxylate metabolism ($P$ = 0.00429), 5. arginine biosynthesis ($P$ = 0.01473), 6. alanine, aspartate and glutamate metabolism ($P$ = 0.02019), 7. glycerolipid metabolism ($P$ = 0.02019), 8. carbon fixation in photosynthetic organisms ($P$ = 0.02217), 9. galactose metabolism ($P$ = 0.03566), 10. amino sugar and nucleotide sugar metabolism ($P$ = 0.07616), 11. fructose and mannose metabolism ($P$ = 0.13874), 12. tyrosine metabolism ($P$ = 0.1872), 13. phenylalanine, tyrosine and tryptophan biosynthesis ($P$ = 0.22414), 14. inositol phosphate metabolism ($P$ = 0.24202), 15. glycolysis/gluconeogenesis ($P$ = 0.24202), 16. phosphatidylinositol signaling system ($P$ = 0.25082), 17. glycine, serine and threonine metabolism ($P$ = 0.27665), 18. glycerophospholipid metabolism ($P$ = 0.34157); detailed results are presented in Supplemental Table 2.

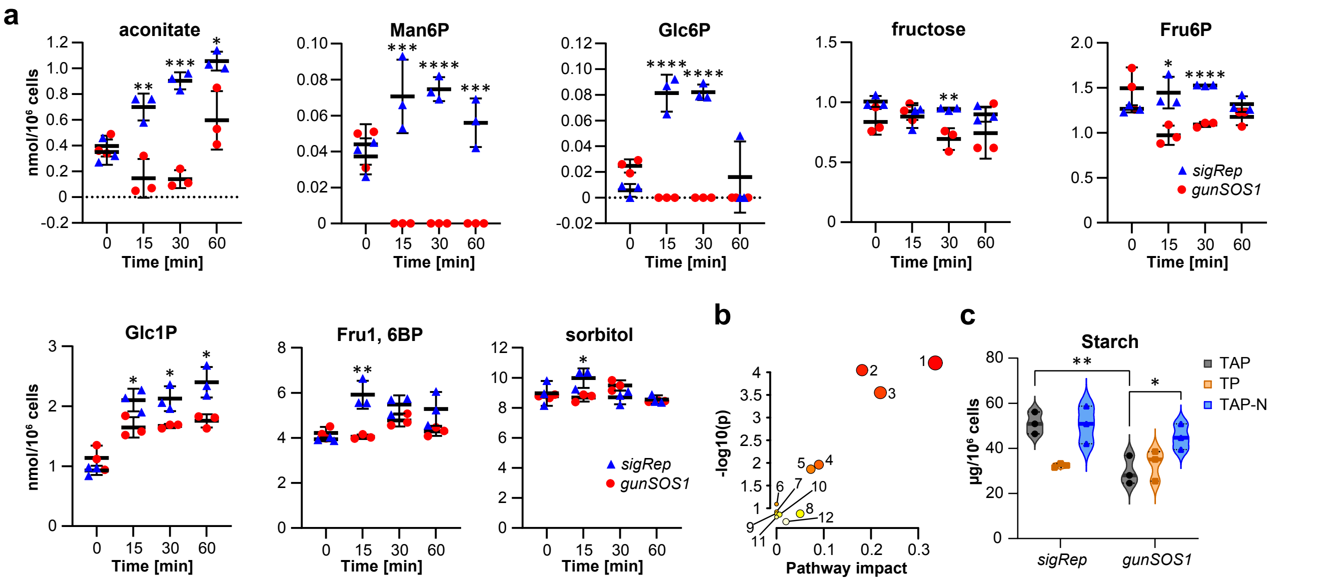


**Supplementary Fig. 6 Metabolites with significantly decreased content in *gunSOS1* compared to *sigRep* in the light. a** LC-MS/MS analysis; cis-aconitic acid (aconitate), mannose 6-phosphate (Man6P), glucose 6-phosphate (Glc6P), D-fructose (fructose), fructose 6-phosphate (Fru6P), glucose 1-phosphate (Glc1P), fructose 1,6-bisphosphate (Fru1, 6BP), and sorbitol. Measurements were performed in biological triplicates ($n$ = 3), horizontal bars represent the calculated mean, vertical error bars represent calculated ±SD; significant differences were calculated using two-tailed Student's $t$-test and are indicated by asterisks (non-significant not shown), ***P** < 0.05, **$P$ < 0.01, ***$P$ < 0.001, and ****$P$ < 0.0001. **b** Impact of deficient metabolites on cell metabolism analysed using MetaboAnalyst 5.0 (https://www.metaboanalyst.ca), sorted by the $P$ value; 1. fructose and mannose metabolism ($P$ = 6.23E-05), 2. amino sugar and nucleotide sugar metabolism ($P$ = 8.99E-05), 3. starch and sucrose metabolism ($P$ = 0.000282), 4. Glycolysis / Gluconeogenesis ($P$ = 0.01094), 5. Galactose metabolism ($P$ = 0.013767), 6. pentose and glucuronate interconversions ($P$ = 0.081139), 7. pentose phosphate pathway ($P$ = 0.1195), 8. tricarboxylic acid cycle (TCA cycle; $P$ = 0.13197), 9. glycerolipid metabolism ($P$ = 0.13197), 10. carbon fixation in photosynthetic organisms ($P$ = 0.13815), 11. inositol phosphate metabolism ($P$ = 0.15646), 12. glyoxylate and dicarboxylate metabolism ($P$ = 0.19785); detailed results are presented in Supplemental Table 3. **c** Starch accumulation in *gunSOS1* compared to *sigRep*. Measurements were performed in biological triplicates ($n$ = 3); median is shown as a center line; upper and lower quartiles are shown as dotted lines. Significant differences were calculated using two-tailed Student's $t$-test and are indicated by asterisks (non-significant not shown), *$P$ < 0.05, **$P$ < 0.01.

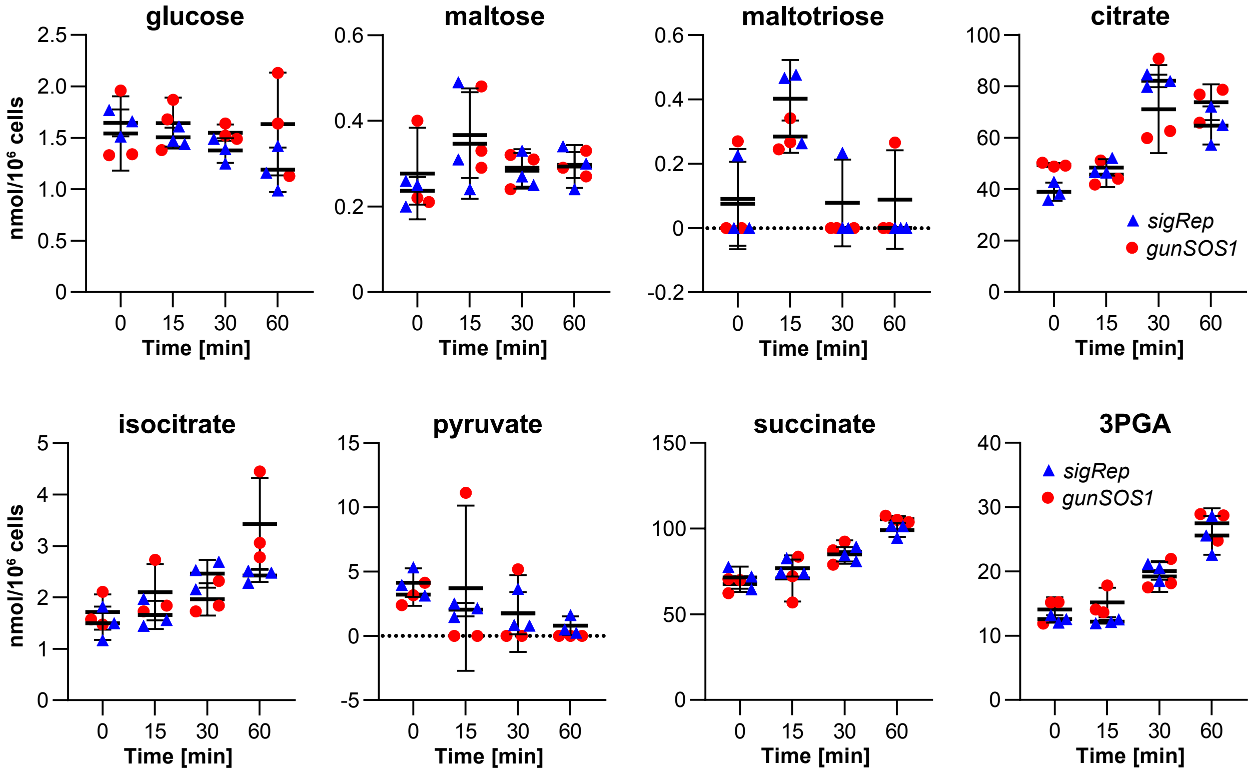


**Supplementary Fig. 7 Metabolites content in *gunSOS1* compared to *sigRep*.** D-glucose (glucose), D-maltose (maltose), maltotriose, citric acid (citrate), isocitric acid (isocitrate), pyruvic acid (pyruvate), succinic acid (succinate), 3-phosphoglyceric acid (3PGA). Measurements were performed in biological triplicates and are presented as mean ($n$ = 3), horizontal bars represent the calculated mean, vertical error bars represent calculated ±SD; significant differences were calculated using two-tailed Student's $t$-test; metabolites presented here did not show significant change in light.

a

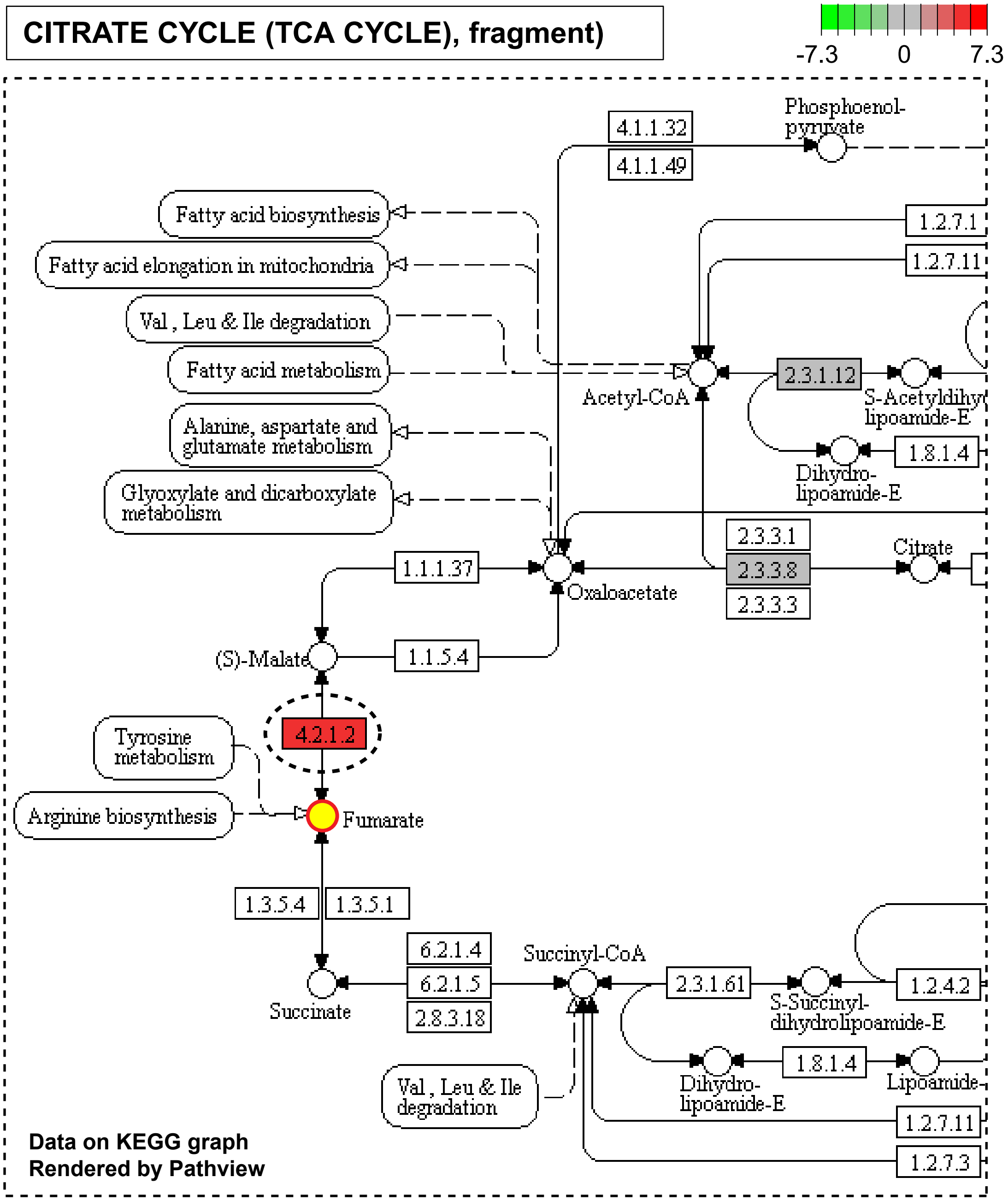

CITRATE CYCLE (TCA CYCLE), fragment)
-7.3
0
7.3
Phosphoenol-pyruvate
4.1.1.32
4.1.1.49
Fatty acid biosynthesis
Fatty acid elongation in mitochondria
Val , Leu & Ile degradation
Fatty acid metabolism
1.2.7.1
1.2.7.11
Acetyl-CoA
2.3.1.12
S-Acetyldihy
lipoamide-E
1.8.1.4
Dihydro-lipoamide-E
Alanine, aspartate and glutamate metabolism
Glyoxylate and dicarboxylate metabolism
2.3.3.1
2.3.3.8
2.3.3.3
Citrate
1.1.1.37
Oxaloacetate
(S)-Malate
1.1.5.4
4.2.1.2
Tyrosine metabolism
Arginine biosynthesis
Fumarate
1.3.5.4
1.3.5.1
6.2.1.4
6.2.1.5
2.8.3.18
Succinyl-CoA
2.3.1.61
S-Succinyl-dihydrolipoamide-E
1.2.4.2
Succinate
1.8.1.4
Dihydro-lipoamide-E
Lipoamide-
Val , Leu & Ile degradation
1.2.7.11
1.2.7.3
Data on KEGG graph
Rendered by Pathview

b

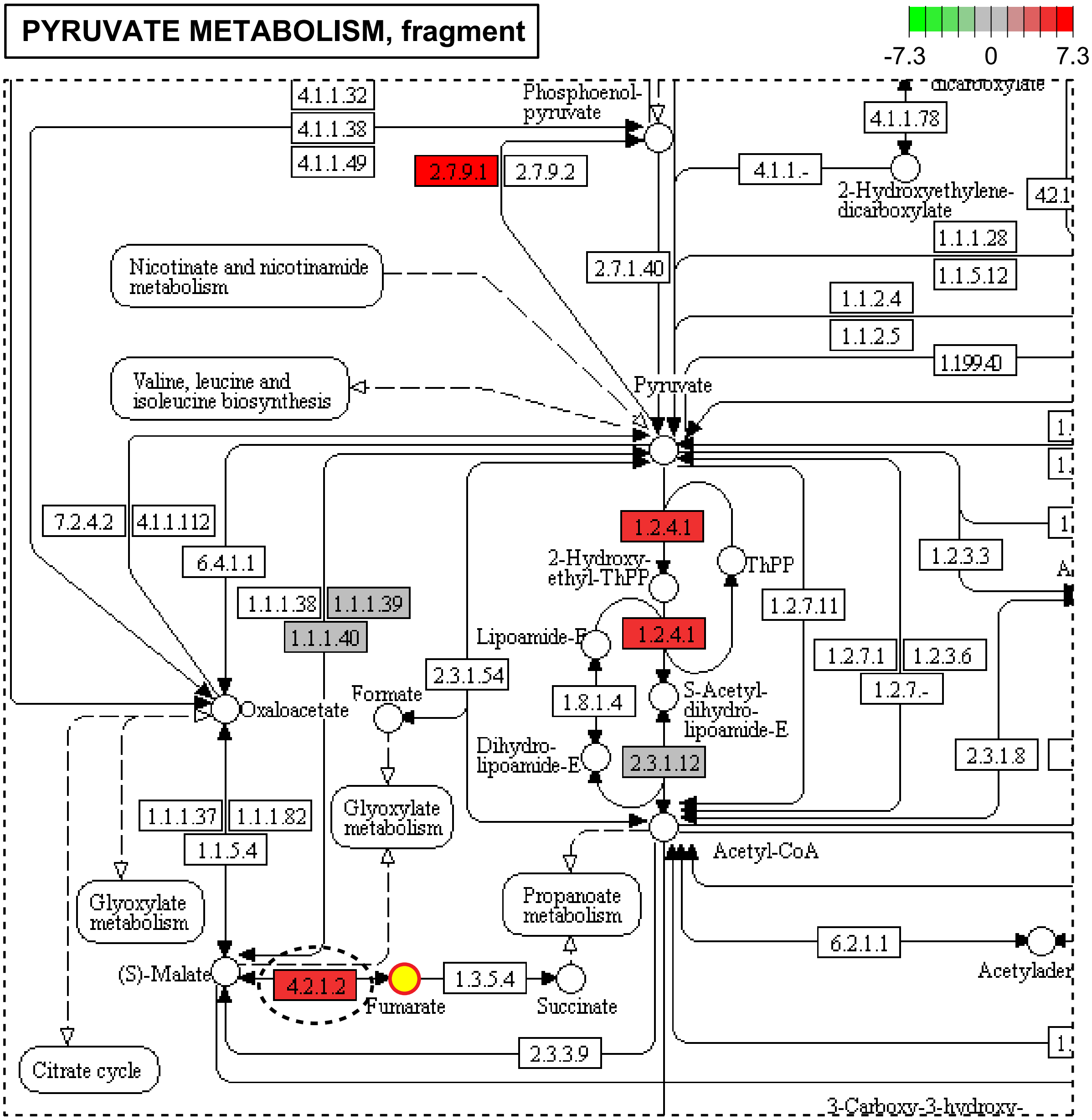


Data on KEGG graph
Rendered by Pathview

**c**

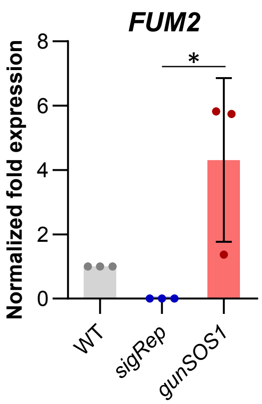


**d**

| EC 4.2.1.2 | FUM1/FUM2 |
| --- | --- |
| Name | fumarate hydratase;<br>fumarase;<br>L-malate hydro-lyase;<br>(S)-malate hydro-lyase |
| Class | Lyases;<br>Carbon-oxygen lyases;<br>Hydro-lyases |
| Sysname | (S)-malate hydro-lyase (fumarate-forming) |
| Reaction(IUBMB) | (S)-malate = fumarate + $H_2O$ [RN:R01082] |
| Reaction(KEGG) | R01082 |
| Substrate | (S)-malate [CPD:C00149] |
| Product | **fumarate** [CPD:C00122];<br>$H_2O$ [CPD:C00001] |

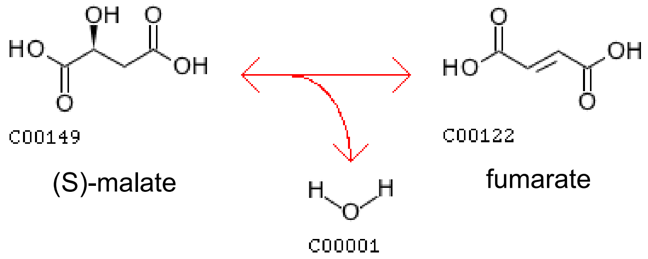


**Supplementary Fig. 8 Accumulation of fumarate in *gunSOS1* compared to *sigRep* is explained by increased expression of *FUMARATE HYDRATASE 2*. a** Fragment of the citrate cycle (TCA cycle) with indicated fumarate. **b** Fragment of the pyruvate metabolism with indicated fumarate. The RNA-seq results showed 5.17-fold increase in *FUMARATE HYDRATASE 2* (*FUM2*, Cre01.g020223.v5.5, EC 4.2.1.2, marked with dashed-line oval) expression in *gunSOS1* compared to *sigRep*. **c** qRT-PCR showed 4.3 times higher *FUM2* transcript in *gunSOS1* compared to WT, while it was not detectable in *sigRep*. Results are presented as normalized fold expression ($2^{-\Delta\Delta Ct}$, WT = 1); experiment was performed in biological replications ($n = 3$); the error bars represent calculated ±SD. Significant differences were calculated using one-way ANOVA, pair-wise comparison with the Tukey's post-hoc test (non-significant not shown), *$P < 0.05$. **d** FUM2 (EC 4.2.1.2) catalyses hydrolysis of malate to fumarate and $H_2O$, based on KEGG analysis.

a

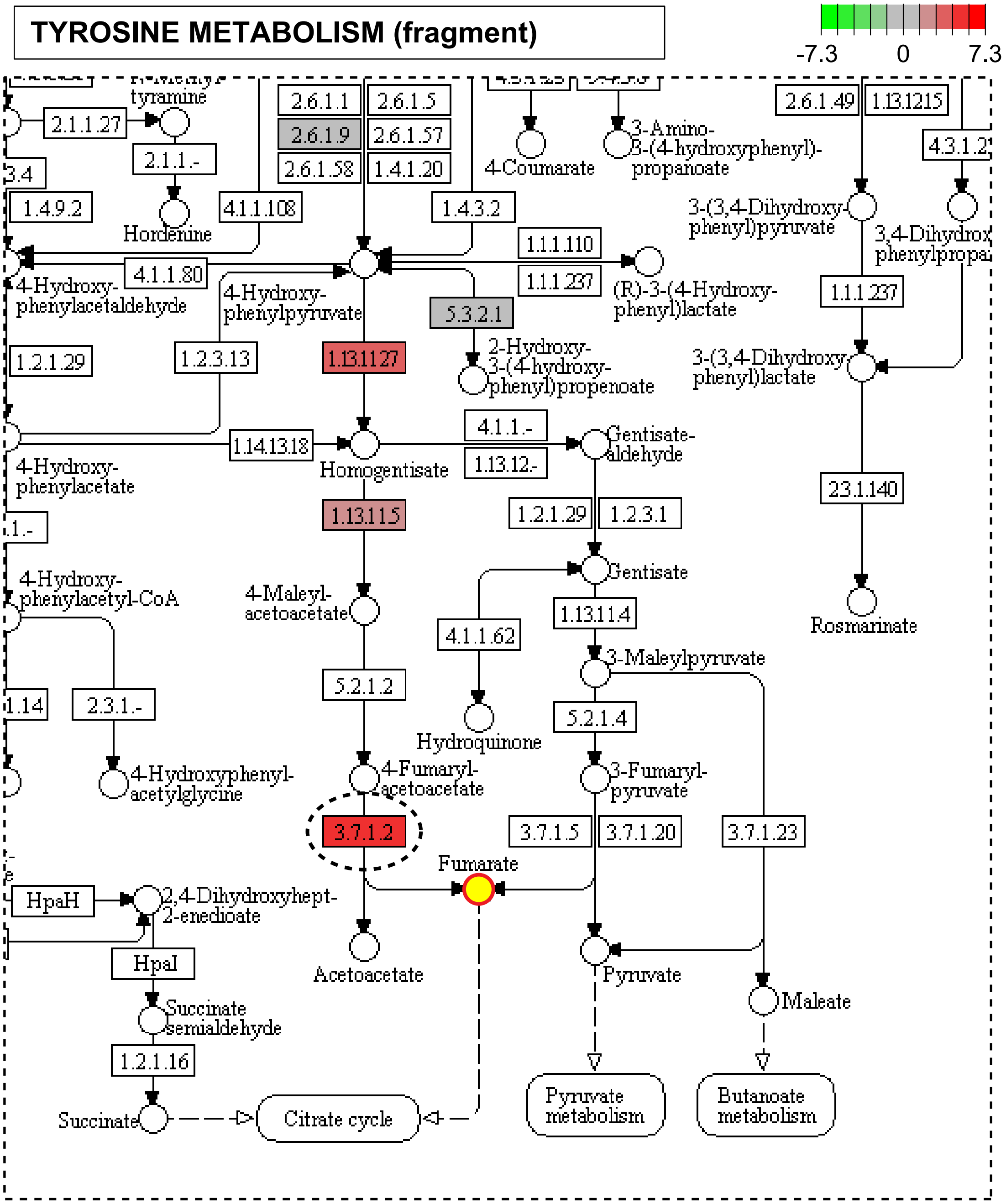


**Data on KEGG graph**
**Rendered by Pathview**

**b**

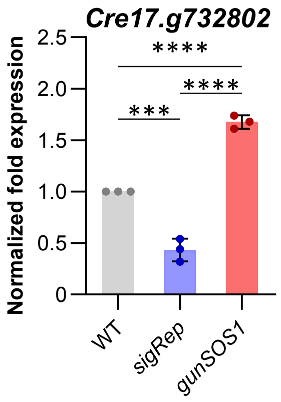


**c**

| EC 3.7.1.2 | Cre17.g732802.v5.5 |
|---|---|
| Name | fumarylacetoacetase; beta-diketonase; fumarylacetoacetate hydrolase |
| Class | Acting on carbon-carbon bonds; In ketonic substances |
| Sysname | 4-fumarylacetoacetate fumarylhydrolase |
| Reaction(IUBMB) | 4-fumarylacetoacetate + $H_2O$ = acetoacetate + fumarate [RN:R01364] |
| Reaction(KEGG) | R01364 |
| Substrate | 4-fumarylacetoacetate [CPD:C01061]; $H_2O$ [CPD:C00001] |
| Product | acetoacetate [CPD:C00164];<br>fumarate [CPD:C00122] |

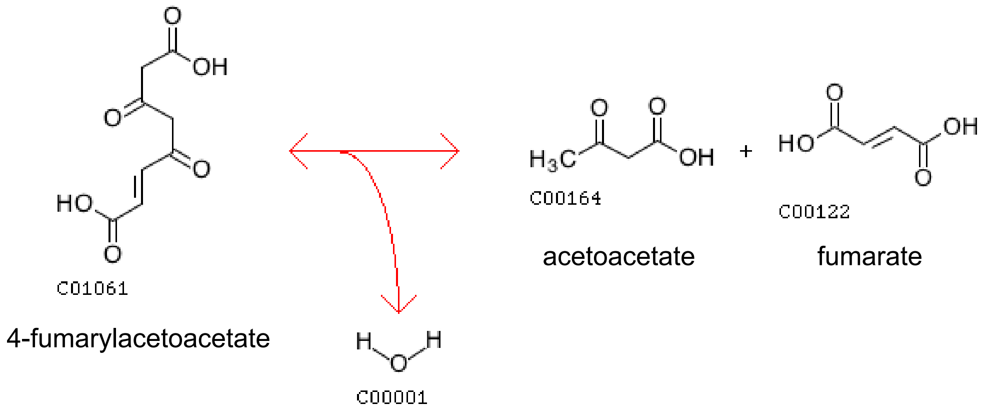


**Supplementary Fig. 9 Accumulation of fumarate in *gunSOS1* compared to *sigRep* is explained by increased expression of *FUMARYLACETOACETASE*. a** Fragment of the tyrosine metabolism with indicated fumarate, based on KEGG, modified. The RNA-seq results showed 4.71-fold increase in *FUMARYLACETOACETASE* (Cre17.g732802, EC 3.7.1.2, marked with dashed-line oval) expression in *gunSOS1* compared to *sigRep*. **b** qRT-PCR showed 3.88-fold increase in *FUMARYLACETOACETASE* (Cre17.g732802) transcript in *gunSOS1* compared to *sigRep*. Results are presented as normalized fold expression ($2^{-\Delta\Delta Ct}$, WT = 1); experiment was performed in biological replications ($n$ = 3); the error bars represent calculated ±SD. Significant differences were calculated using one-way ANOVA, pair-wise comparison with the Tukey's post-hoc test, ****P* < 0.001 and *****P* < 0.0001. **c** FUMARYLACETOACETASE (EC 3.7.1.2) catalyses hydrolysis of fumarylacetoacetate producing acetoacetate, fumarate, and $H_2O$, based on KEGG analysis.

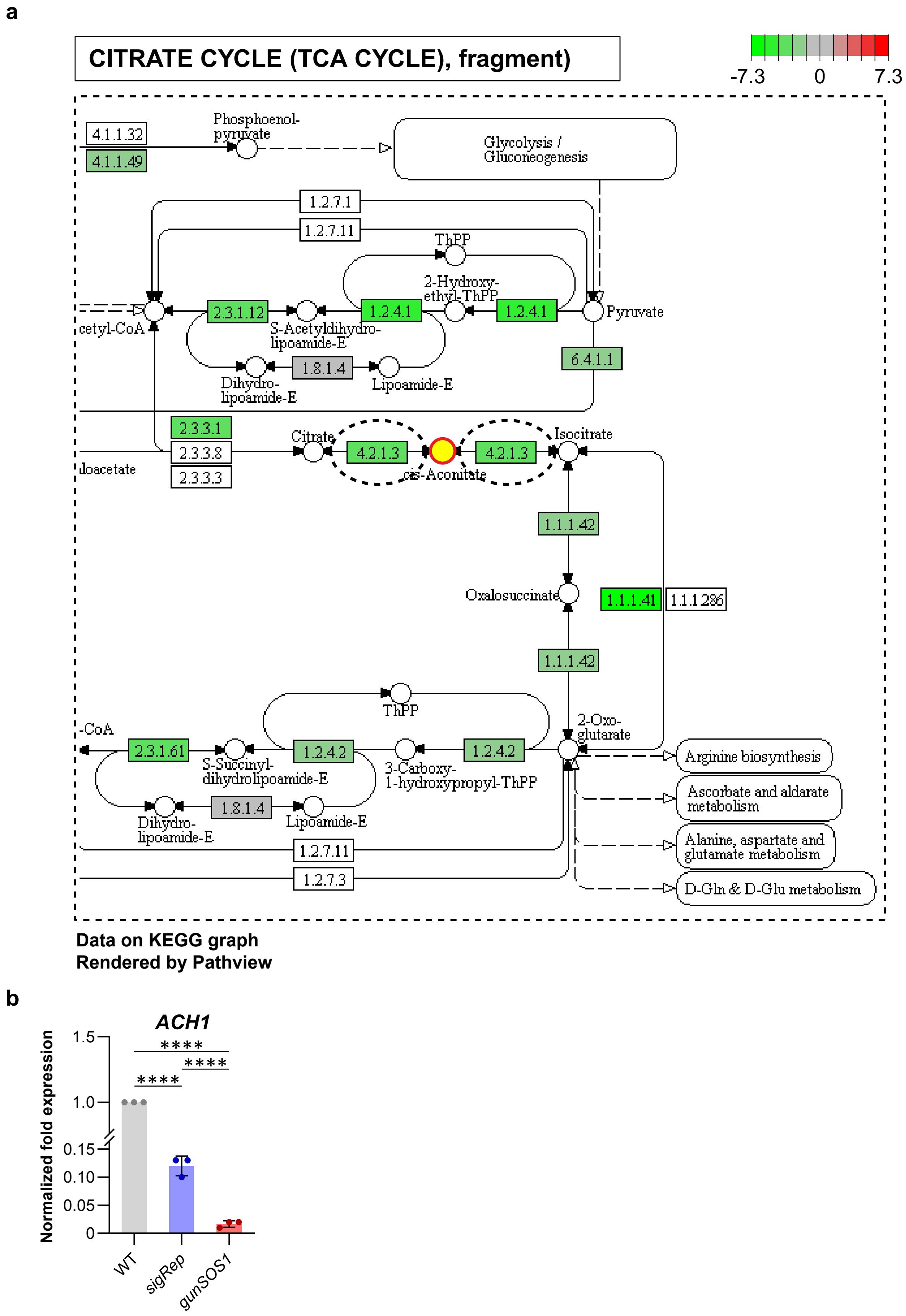
a
CITRATE CYCLE (TCA CYCLE), fragment)
-7.3
0
7.3
4.1.1.32
4.1.1.49
Phosphoenol-
pyruvate
Glycolysis /
Gluconeogenesis
1.2.7.1
1.2.7.11
ThPP
2-Hydroxy-
ethyl-ThPP
2.3.1.12
1.2.4.1
1.2.4.1
Pyruvate
cetyl-CoA
S-Acetyldihydro-
lipoamide-E
1.8.1.4
Dihydro-
lipoamide-E
Lipoamide-E
6.4.1.1
2.3.3.1
2.3.3.8
2.3.3.3
Citrate
4.2.1.3
4.2.1.3
Isocitrate
cis-Aconitate
loacetate
1.1.1.42
Oxalosuccinate
1.1.1.41
1.1.1.286
1.1.1.42
ThPP
-CoA
2.3.1.61
1.2.4.2
1.2.4.2
2-Oxo-
glutarate
S-Succinyl-
dihydrolipoamide-E
3-Carboxy-
1-hydroxypropyl-ThPP
1.8.1.4
Dihydro-
lipoamide-E
Lipoamide-E
1.2.7.11
1.2.7.3
Arginine biosynthesis
Ascorbate and aldarate
metabolism
Alanine, aspartate and
glutamate metabolism
D-Gln & D-Glu metabolism
Data on KEGG graph
Rendered by Pathview
b
ACH1
Normalized fold expression
1.5
1.0
0.15
0.10
0.05
0
****
****
****
WT
sigRep
gunSOS1

**c**

| EC 4.2.1.3 | ACH1 |
|---|---|
| Name | aconitate hydratase; cis-aconitase; aconitase; AcnB; 2-methylaconitate hydratase; citrate(isocitrate) hydro-lyase |
| Class | Lyases; Carbon-oxygen lyases; Hydro-lyases |
| Sysname | citrate(isocitrate) hydro-lyase (cis-aconitate-forming) |
| Reaction(IUBMB) | citrate = isocitrate (overall reaction) [RN:R01324];<br>(1a) citrate = cis-aconitate + $H_2O$ [RN:R01325];<br>(1b) cis-aconitate + $H_2O$ = isocitrate [RN:R01900] |
| Reaction(KEGG) | R01324 R01325 R01900 |
| Substrate | citrate [CPD:C00158]; cis-aconitate [CPD:C00417]; $H_2O$ [CPD:C00001] |
| Product | isocitrate [CPD:C00311]; cis-aconitate [CPD:C00417]; $H_2O$ [CPD:C00001] |
| Comment | Besides interconverting citrate and cis-aconitate, it also interconverts cis-aconitate with isocitrate and, hence, interconverts citrate and isocitrate. The equilibrium mixture is 91% citrate, 6% isocitrate and 3% aconitate. cis-aconitate is used to designate the isomer (Z)-prop-1-ene-1,2,3-tricarboxylate. An iron-sulfur protein, containing a [4Fe-4S] cluster to which the substrate binds. |

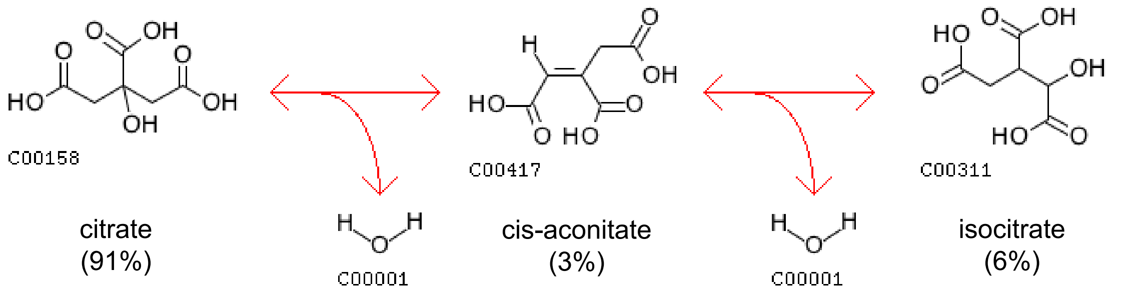


**Supplementary Fig. 10 Deficiency in aconitate in *gunSOS1* compared to *sigRep* is explained by decreased expression of *ACONITATE HYDRATASE 1*. a** Fragment of the TCA cycle with indicated aconitate, based on KEGG, modified. The RNA-seq results showed 2.81-fold decrease in *ACONITATE HYDRATASE 1* (*ACH1*; Cre01.g042750; EC 4.2.1.3) expression in *gunSOS1* compared to *sigRep*. **b** Based on qRT-PCR, *ACH1* transcript level was neraly 8 times lower in *gunSOS1* compared to *sigRep*. Results are presented as normalized fold expression ($2^{-\Delta\Delta Ct}$, WT = 1); experiment was performed in biological replications ($n$ = 3); the error bars represent calculated ±SD. Significant differences were calculated using one-way ANOVA, pair-wise comparison with the Tukey's post-hoc test, ****$P$ < 0.0001. **c** Catalytic reaction of ACH1 (EC 4.2.1.3), based on KEGG analysis.

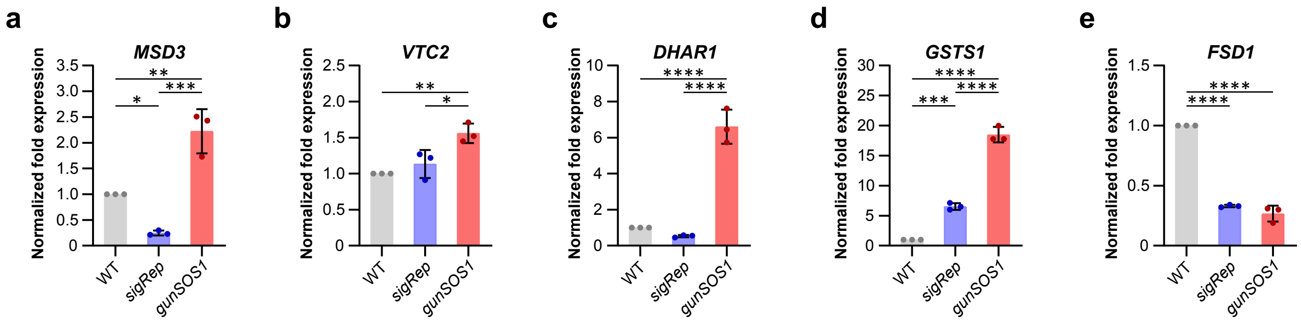


**Supplementary Fig. 11 Expression of genes, which induction was previously associated with $H_2O_2$ or organic peroxides, but not with $^1O_2$. a** *Mn SUPEROXIDE DISMUTASE 3* (*MSD3*). **b** GDP-L-galactose *PHOSPHORYLASE* (*VTC2*). **c** *DEHYDROASCORBATE REDUCTASE* (*DHAR1*). **d** *GLUTATHIONE S-TRANSFERASE* (*GSTS1*). **e** Fe *SUPEROXIDE DISMUTASE* (*FSD1*). Experiments were performed in biological replications ($n$ = 3); results are presented as normalized fold expression ($2^{-\Delta\Delta Ct}$, WT = 1); the error bars represent calculated ±SD. Significant differences were calculated using one-way ANOVA, pair-wise comparison with the Tukey's post-hoc test (non-significant not shown), *$P$ < 0.05, **$P$ < 0.01, ***$P$ < 0.001, and ****$P$ < 0.0001.
