## Supplementary material for "Singlet oxygen-induced signalling depends on the metabolic status of the Chlamydomonas cell": Al Youssef et al_Supplementary Tables

**Supplementary Table 1. Metabolites analyzed in *gunSOS1* compared to *sigRep* (LC-MS/MS).** Names and annotations are based on the MetaboAnalyst5.0 portal (https://www.metaboanalyst.ca); The Human Metabolome Database (HMDB), Kyoto Encyclopedia of Genes and Genomes (KEGG).

| **Sugars** | | |
|---|---|---|
| **Match** | **HMDB** | **KEGG** |
| Trehalose 6-phosphate | HMDB0001124 | C00689 |
| Trehalose | HMDB0000975 | C01083 |
| D-Glucose | HMDB0000122 | C00031 |
| D-Fructose | HMDB0000660 | C02336 |
| D-Maltose | HMDB0000163 | C00208 |
| Maltotriose | HMDB0001262 | C01835 |
| ADP-glucose | HMDB0006557 | C00498 |
| Galactose 1-phosphate | HMDB0000645 | C00446 |
| Fructose 1-phosphate | HMDB0001076 | C01094 |
| Alpha-D-Glucose 1,6-bisphosphate | HMDB0003514 | C01231 |
| Glucose 1-phosphate | HMDB0001586 | C00446 |
| Fructose 6-phosphate | HMDB0000124 | C00085 |
| Mannose 6-phosphate | HMDB0001078 | C00275 |
| Fructose 1,6-bisphosphate | HMDB0001058 | C00354 |
| Glucose 6-phosphate | HMDB0001401 | C00092 |
| **Sugar alcohols and non-sugar metabolites** | | |
| **Match** | **HMDB** | **KEGG** |
| Sorbitol | HMDB0000247 | C00794 |
| myo-Inositol | HMDB0000211 | C00137 |
| Glycerol 3-phosphate | HMDB0000126 | C00093 |
| Phosphoenolpyruvic acid | HMDB0000263 | C00074 |
| cis-Aconitic acid | HMDB0000072 | C00417 |
| Isocitric acid | HMDB0000193 | C00311 |
| Oxoglutaric acid | HMDB0000208 | C00026 |
| Pyruvic acid | HMDB0000243 | C00022 |
| Succinic acid | HMDB0000254 | C00042 |
| 3-Phosphoglyceric acid | HMDB0000807 | C00597 |
| Glyceric acid | HMDB0000139 | C00258 |
| Citric acid | HMDB0000094 | C00158 |
| L-Malic acid | HMDB0000156 | C00149 |
| Fumaric acid | HMDB0000134 | C00122 |

**Supplementary Table 2. Impact of accumulating metabolites on cell metabolism.** Details for the analysis performed using MetaboAnalyst 5.0 portal (https://www.metaboanalyst.ca), presented in Supplementary Fig. 5b; sorted by the *P* value.

| Pathway name | Total | Expected | Hits | *P* | -log(*P*) | Holm *P* | FDR | Impact |
|---|---|---|---|---|---|---|---|---|
| Citrate cycle (TCA cycle) | 20 | 0.22727 | 4 | $4.41 \times 10^{-5}$ | 4.3558 | 0.0037 | 0.0037 | 0.132 |
| Pyruvate metabolism | 21 | 0.23864 | 3 | 0.00136 | 2.8674 | 0.11263 | 0.038 | 0.042 |
| Starch and sucrose metabolism | 21 | 0.23864 | 3 | 0.00136 | 2.8674 | 0.11263 | 0.038 | 0.032 |
| Glyoxylate and dicarboxylate metabolism | 31 | 0.35227 | 3 | 0.00429 | 2.3674 | 0.34764 | 0.09013 | 0.118 |
| Arginine biosynthesis | 17 | 0.19318 | 2 | 0.01473 | 1.8317 | 1 | 0.2328 | 0 |
| Alanine, aspartate and glutamate metabolism | 20 | 0.22727 | 2 | 0.02019 | 1.6949 | 1 | 0.2328 | 0.098 |
| Glycerolipid metabolism | 20 | 0.22727 | 2 | 0.02019 | 1.6949 | 1 | 0.2328 | 0.045 |
| Carbon fixation in photosynthetic organisms | 21 | 0.23864 | 2 | 0.02217 | 1.6542 | 1 | 0.2328 | 0.063 |
| Galactose metabolism | 27 | 0.30682 | 2 | 0.03566 | 1.4478 | 1 | 0.33281 | 0 |
| Amino sugar and nucleotide sugar metabolism | 41 | 0.46591 | 2 | 0.07616 | 1.1183 | 1 | 0.63976 | 0 |
| Fructose and mannose metabolism | 13 | 0.14773 | 1 | 0.13874 | 0.85779 | 1 | 1 | 0.004 |
| Tyrosine metabolism | 18 | 0.20455 | 1 | 0.1872 | 0.7277 | 1 | 1 | 0 |
| Phenylalanine, tyrosine and tryptophan biosynthesis | 22 | 0.25 | 1 | 0.22414 | 0.64948 | 1 | 1 | 0 |
| Inositol phosphate metabolism | 24 | 0.27273 | 1 | 0.24202 | 0.61615 | 1 | 1 | 0.205 |
| Glycolysis / Gluconeogenesis | 24 | 0.27273 | 1 | 0.24202 | 0.61615 | 1 | 1 | 0.125 |
| Phosphatidylinositol signaling system | 25 | 0.28409 | 1 | 0.25082 | 0.60064 | 1 | 1 | 0.185 |
| Glycine, serine and threonine metabolism | 28 | 0.31818 | 1 | 0.27665 | 0.55807 | 1 | 1 | 0.041 |
| Glycerophospholipid metabolism | 36 | 0.40909 | 1 | 0.34157 | 0.46652 | 1 | 1 | 0.099 |

**Supplementary Table 3. Impact of deficient metabolites on cell metabolism**. Details for the analysis performed using MetaboAnalyst 5.0 portal (https://www.metaboanalyst.ca), presented in Supplementary Fig. 6b; sorted by the *P* value.

| Pathway name | Total | Expected | Hits | *P* | -log(*P*) | Holm *P* | FDR | Impact |
|---|---|---|---|---|---|---|---|---|
| Fructose and mannose metabolism | 13 | 0.090909 | 3 | $6.23 \times 10^{-5}$ | 4.2058 | 0.00523 | 0.003775 | 0.336 |
| Amino sugar and nucleotide sugar metabolism | 41 | 0.28671 | 4 | $8.99 \times 10^{-5}$ | 4.0463 | 0.007461 | 0.003775 | 0.181 |
| Starch and sucrose metabolism | 21 | 0.14685 | 3 | 0.000282 | 3.5498 | 0.023122 | 0.007895 | 0.22 |
| Glycolysis / Gluconeogenesis | 24 | 0.16783 | 2 | 0.01094 | 1.961 | 0.88616 | 0.22974 | 0.09 |
| Galactose metabolism | 27 | 0.18881 | 2 | 0.013767 | 1.8612 | 1 | 0.23128 | 0.073 |
| Pentose and glucuronate interconversions | 12 | 0.083916 | 1 | 0.081139 | 1.0908 | 1 | 1 | 0 |
| Pentose phosphate pathway | 18 | 0.12587 | 1 | 0.1195 | 0.92262 | 1 | 1 | 0 |
| Citrate cycle (TCA cycle) | 20 | 0.13986 | 1 | 0.13197 | 0.87951 | 1 | 1 | 0.05 |
| Glycerolipid metabolism | 20 | 0.13986 | 1 | 0.13197 | 0.87951 | 1 | 1 | 0 |
| Carbon fixation in photosynthetic organisms | 21 | 0.14685 | 1 | 0.13815 | 0.85964 | 1 | 1 | 0.006 |
| Inositol phosphate metabolism | 24 | 0.16783 | 1 | 0.15646 | 0.8056 | 1 | 1 | 0 |
| Glyoxylate and dicarboxylate metabolism | 31 | 0.21678 | 1 | 0.19785 | 0.70366 | 1 | 1 | 0.02 |

**Supplementary Table 4. Statistical analyses for $\mathit{GPX5}_{cyt}$ expression following treatment with fumarate, presented in Fig. 6a.** *P*, probability (significance test); *SS*, sum of squares; *DF*, degree of freedom; *MS*, means square; *F*, F-value (Fisher test); *DFn*, degrees of freedom in the numerator; *DFd*, degrees of freedom in the denominator; *SE*, standard error of the sample mean; *N*, sample number.

| **Two-way ANOVA Ordinary** | | | | | |
|---|---|---|---|---|---|
| Alpha | 0.05 | | | | |
| **Source of Variation** | **% of total variation** | **P value** | **P value summary** | **Significant?** | |
| Interaction | 27.86 | <0.0001 | **** | Yes | |
| strain | 54.47 | <0.0001 | **** | Yes | |
| metabolite | 15.36 | <0.0001 | **** | Yes | |
| **ANOVA table** | **SS** | **DF** | **MS** | **F (DFn, DFd)** | **P value** |
| Interaction | 120.7 | 6 | 20.11 | F (6, 24) = 48.21 | P<0.0001 |
| strain | 236.0 | 2 | 118.0 | F (2, 24) = 282.8 | P<0.0001 |
| metabolite | 66.51 | 3 | 22.17 | F (3, 24) = 53.15 | P<0.0001 |
| Residual | 10.01 | 24 | 0.4172 | | |

| **Data summary** | |
|---|---|
| Number of columns (metabolite) | 4 |
| Number of rows (strain) | 3 |
| Number of values | 36 |

| **Two-way ANOVA Multiple comparison** | |
|---|---|
| Number of families | 3 |
| Number of comparisons per family | 3 |
| Alpha | 0.05 |

| **Dunnett's multiple comparisons test** | **Mean Diff.** | **95.00% CI of diff.** | **Below threshold?** | **Summary** | **Adjusted P Value** |
|---|---|---|---|---|---|
| WT | | | | | |
| control vs. 20 µM Fum | 0.7367 | -0.5854 to 2.059 | No | ns | 0.3810 |
| control vs. 50 µM Fum | 0.6733 | -0.6487 to 1.995 | No | ns | 0.4509 |
| control vs. 100 µM Fum | 0.6733 | -0.6487 to 1.995 | No | ns | 0.4509 |
| *sigRep* | | | | | |
| control vs. 20 µM Fum | 4.690 | 3.368 to 6.012 | Yes | **** | <0.0001 |
| control vs. 50 µM Fum | 7.970 | 6.648 to 9.292 | Yes | **** | <0.0001 |
| control vs. 100 µM Fum | 10.53 | 9.211 to 11.86 | Yes | **** | <0.0001 |
| *gunSOS1* | | | | | |
| control vs. 20 µM Fum | -0.06667 | -1.389 to 1.255 | No | ns | 0.9985 |
| control vs. 50 µM Fum | -0.2433 | -1.565 to 1.079 | No | ns | 0.9387 |
| control vs. 100 µM Fum | -0.2600 | -1.582 to 1.062 | No | ns | 0.9270 |

| **Test details** | **Mean 1** | **Mean 2** | **Mean Diff.** | **SE of diff.** | **N1** | **N2** | **q** | **DF** |
|---|---|---|---|---|---|---|---|---|
| WT | | | | | | | | |
| control vs. 20 µM Fum | 1.000 | 0.2633 | 0.7367 | 0.5274 | 3 | 3 | 1.397 | 24.00 |
| control vs. 50 µM Fum | 1.000 | 0.3267 | 0.6733 | 0.5274 | 3 | 3 | 1.277 | 24.00 |
| control vs. 100 µM Fum | 1.000 | 0.3267 | 0.6733 | 0.5274 | 3 | 3 | 1.277 | 24.00 |
| *sigRep* | | | | | | | | |
| control vs. 20 µM Fum | 11.59 | 6.903 | 4.690 | 0.5274 | 3 | 3 | 8.893 | 24.00 |
| control vs. 50 µM Fum | 11.59 | 3.623 | 7.970 | 0.5274 | 3 | 3 | 15.11 | 24.00 |
| control vs. 100 µM Fum | 11.59 | 1.060 | 10.53 | 0.5274 | 3 | 3 | 19.97 | 24.00 |
| *gunSOS1* | | | | | | | | |
| control vs. 20 µM Fum | 0.1133 | 0.1800 | -0.06667 | 0.5274 | 3 | 3 | 0.1264 | 24.00 |
| control vs. 50 µM Fum | 0.1133 | 0.3567 | -0.2433 | 0.5274 | 3 | 3 | 0.4614 | 24.00 |
| control vs. 100 µM Fum | 0.1133 | 0.3733 | -0.2600 | 0.5274 | 3 | 3 | 0.4930 | 24.00 |

**Supplementary Table 5. Statistical analyses for *GPX5$_{cp}$* expression following treatment with fumarate, presented in Fig. 6a.** *P*, probability (significance test); *SS*, sum of squares; *DF*, degree of freedom; *MS*, means square; *F*, F-value (Fisher test); *DFn*, degrees of freedom in the numerator; *DFd*, degrees of freedom in the denominator; *SE*, standard error of the sample mean; *N*, sample number.

| **Two-way ANOVA Ordinary** | | | | | |
|---|---|---|---|---|---|
| Alpha | 0.05 | | | | |
| **Source of Variation** | **% of total variation** | **P value** | **P value summary** | **Significant?** | |
| Interaction | 34.57 | <0.0001 | **** | Yes | |
| strain | 40.98 | <0.0001 | **** | Yes | |
| metabolite | 23.29 | <0.0001 | **** | Yes | |
| **ANOVA table** | **SS** | **DF** | **MS** | **F (DFn, DFd)** | **P value** |
| Interaction | 19.99 | 6 | 3.332 | F (6, 24) = 118.9 | P<0.0001 |
| strain | 23.69 | 2 | 11.85 | F (2, 24) = 422.6 | P<0.0001 |
| metabolite | 13.47 | 3 | 4.489 | F (3, 24) = 160.1 | P<0.0001 |
| Residual | 0.6727 | 24 | 0.02803 | | |
| **Data summary** | | | | | |
| Number of columns (metabolite) | 4 | | | | |
| Number of rows (strain) | 3 | | | | |
| Number of values | 36 | | | | |

| **Two-way ANOVA Multiple comparison** | | | | | |
|---|---|---|---|---|---|
| Number of families | 3 | | | | |
| Number of comparisons per family | 3 | | | | |
| Alpha | 0.05 | | | | |
| **Dunnett's multiple comparisons test** | **Mean Diff.** | **95.00% CI of diff.** | **Below threshold?** | **Summary** | **Adjusted P Value** |
| WT | | | | | |
| control vs. 20 µM Fum | 0.7500 | 0.4073 to 1.093 | Yes | **** | <0.0001 |
| control vs. 50 µM Fum | 0.6767 | 0.3340 to 1.019 | Yes | *** | 0.0001 |
| control vs. 100 µM Fum | 0.7033 | 0.3606 to 1.046 | Yes | **** | <0.0001 |
| *sigRep* | | | | | |
| control vs. 20 µM Fum | 1.353 | 1.011 to 1.696 | Yes | **** | <0.0001 |
| control vs. 50 µM Fum | 3.690 | 3.347 to 4.033 | Yes | **** | <0.0001 |
| control vs. 100 µM Fum | 3.930 | 3.587 to 4.273 | Yes | **** | <0.0001 |
| *gunSOS1* | | | | | |
| control vs. 20 µM Fum | -0.02333 | -0.3660 to 0.3194 | No | ns | 0.9965 |
| control vs. 50 µM Fum | -0.1033 | -0.4460 to 0.2394 | No | ns | 0.7919 |
| control vs. 100 µM Fum | -0.1033 | -0.4460 to 0.2394 | No | ns | 0.7919 |

| **Test details** | **Mean 1** | **Mean 2** | **Mean Diff.** | **SE of diff.** | **N1** | **N2** | **q** | **DF** |
|---|---|---|---|---|---|---|---|---|
| WT | | | | | | | | |
| control vs. 20 µM Fum | 1.000 | 0.2500 | 0.7500 | 0.1367 | 3 | 3 | 5.486 | 24.00 |
| control vs. 50 µM Fum | 1.000 | 0.3233 | 0.6767 | 0.1367 | 3 | 3 | 4.950 | 24.00 |
| control vs. 100 µM Fum | 1.000 | 0.2967 | 0.7033 | 0.1367 | 3 | 3 | 5.145 | 24.00 |
| *sigRep* | | | | | | | | |
| control vs. 20 µM Fum | 4.223 | 2.870 | 1.353 | 0.1367 | 3 | 3 | 9.900 | 24.00 |
| control vs. 50 µM Fum | 4.223 | 0.5333 | 3.690 | 0.1367 | 3 | 3 | 26.99 | 24.00 |
| control vs. 100 µM Fum | 4.223 | 0.2933 | 3.930 | 0.1367 | 3 | 3 | 28.75 | 24.00 |
| *gunSOS1* | | | | | | | | |
| control vs. 20 µM Fum | 0.05000 | 0.07333 | -0.02333 | 0.1367 | 3 | 3 | 0.1707 | 24.00 |
| control vs. 50 µM Fum | 0.05000 | 0.1533 | -0.1033 | 0.1367 | 3 | 3 | 0.7559 | 24.00 |
| control vs. 100 µM Fum | 0.05000 | 0.1533 | -0.1033 | 0.1367 | 3 | 3 | 0.7559 | 24.00 |

**Supplementary Table 6. Statistical analyses for *$GPX5_{cyt}$* expression following treatment with aconitate, presented in Fig. 6b.** *P*, probability (significance test); *SS*, sum of squares; *DF*, degree of freedom; *MS*, means square; *F*, F-value (Fisher test); *DFn*, degrees of freedom in the numerator; *DFd*, degrees of freedom in the denominator; *SE*, standard error of the sample mean; *N*, sample number.

| **Two-way ANOVA Ordinary** | | | | | |
|---|---|---|---|---|---|
| Alpha | 0.05 | | | | |
| **Source of Variation** | **% of total variation** | **P value** | **P value summary** | **Significant?** | |
| Interaction | 22.90 | <0.0001 | **** | Yes | |
| strain | 31.91 | <0.0001 | **** | Yes | |
| metabolite | 42.02 | <0.0001 | **** | Yes | |
| **ANOVA table** | **SS** | **DF** | **MS** | **F (DFn, DFd)** | **P value** |
| Interaction | 158.3 | 6 | 26.38 | F (6, 24) = 28.93 | P<0.0001 |
| strain | 220.6 | 2 | 110.3 | F (2, 24) = 120.9 | P<0.0001 |
| metabolite | 290.5 | 3 | 96.83 | F (3, 24) = 106.2 | P<0.0001 |
| Residual | 21.88 | 24 | 0.9119 | | |

| **Data summary** | |
|---|---|
| Number of columns (metabolite) | 4 |
| Number of rows (strain) | 3 |
| Number of values | 36 |

| **Two-way ANOVA Multiple comparison** | | | | | |
|---|---|---|---|---|---|
| Number of families | 3 | | | | |
| Number of comparisons per family | 3 | | | | |
| Alpha | 0.05 | | | | |
| **Dunnett's multiple comparisons test** | **Mean Diff.** | **95.00% CI of diff.** | **Below threshold?** | **Summary** | **Adjusted P Value** |
| WT | | | | | |
| control vs. 20 µM Acon | -0.1767 | -2.131 to 1.778 | No | ns | 0.9918 |
| control vs. 50 µM Acon | -0.6067 | -2.561 to 1.348 | No | ns | 0.7781 |
| control vs. 100 µM Acon | 0.1033 | -1.851 to 2.058 | No | ns | 0.9983 |
| *sigRep* | | | | | |
| control vs. 20 µM Acon | -8.807 | -10.76 to -6.852 | Yes | **** | <0.0001 |
| control vs. 50 µM Acon | 0.6467 | -1.308 to 2.601 | No | ns | 0.7453 |
| control vs. 100 µM Acon | 3.903 | 1.949 to 5.858 | Yes | *** | 0.0001 |
| *gunSOS1* | | | | | |
| control vs. 20 µM Acon | -9.167 | -11.12 to -7.212 | Yes | **** | <0.0001 |
| control vs. 50 µM Acon | -0.3567 | -2.311 to 1.598 | No | ns | 0.9401 |
| control vs. 100 µM Acon | -0.1867 | -2.141 to 1.768 | No | ns | 0.9903 |

| **Test details** | **Mean 1** | **Mean 2** | **Mean Diff.** | **SE of diff.** | **N1** | **N2** | **q** | **DF** |
|---|---|---|---|---|---|---|---|---|
| WT | | | | | | | | |
| control vs. 20 µM Acon | 1.000 | 1.177 | -0.1767 | 0.7797 | 3 | 3 | 0.2266 | 24.00 |
| control vs. 50 µM Acon | 1.000 | 1.607 | -0.6067 | 0.7797 | 3 | 3 | 0.7781 | 24.00 |
| control vs. 100 µM Acon | 1.000 | 0.8967 | 0.1033 | 0.7797 | 3 | 3 | 0.1325 | 24.00 |
| sigRep | | | | | | | | |
| control vs. 20 µM Acon | 6.030 | 14.84 | -8.807 | 0.7797 | 3 | 3 | 11.30 | 24.00 |
| control vs. 50 µM Acon | 6.030 | 5.383 | 0.6467 | 0.7797 | 3 | 3 | 0.8294 | 24.00 |
| control vs. 100 µM Acon | 6.030 | 2.127 | 3.903 | 0.7797 | 3 | 3 | 5.006 | 24.00 |
| gunSOS1 | | | | | | | | |
| control vs. 20 µM Acon | 0.5867 | 9.753 | -9.167 | 0.7797 | 3 | 3 | 11.76 | 24.00 |
| control vs. 50 µM Acon | 0.5867 | 0.9433 | -0.3567 | 0.7797 | 3 | 3 | 0.4574 | 24.00 |
| control vs. 100 µM Acon | 0.5867 | 0.7733 | -0.1867 | 0.7797 | 3 | 3 | 0.2394 | 24.00 |

**Supplementary Table 7. Statistical analyses for *$GPX5_{cp}$* expression following treatment with aconitate, presented in Fig. 6b.** *P*, probability (significance test); *SS*, sum of squares; *DF*, degree of freedom; *MS*, means square; *F*, F-value (Fisher test); *DFn*, degrees of freedom in the numerator; *DFd*, degrees of freedom in the denominator; *SE*, standard error of the sample mean; *N*, sample number.

| **Two-way ANOVA Ordinary** | | | | | |
|---|---|---|---|---|---|
| Alpha | 0.05 | | | | |
| **Source of Variation** | **% of total variation** | **P value** | **P value summary** | **Significant?** | |
| Interaction | 30.75 | <0.0001 | **** | Yes | |
| strain | 27.59 | <0.0001 | **** | Yes | |
| metabolite | 39.86 | <0.0001 | **** | Yes | |
| **ANOVA table** | SS | DF | MS | F (DFn, DFd) | P value |
| Interaction | 31.93 | 6 | 5.321 | F (6, 24) = 68.18 | P<0.0001 |
| strain | 28.65 | 2 | 14.32 | F (2, 24) = 183.5 | P<0.0001 |
| metabolite | 41.40 | 3 | 13.80 | F (3, 24) = 176.8 | P<0.0001 |
| Residual | 1.873 | 24 | 0.07804 | | |
| **Data summary** | | | | | |
| Number of columns (metabolite) | | | | | 4 |
| Number of rows (strain) | | | | | 3 |
| Number of values | | | | | 36 |

| **Two-way ANOVA Multiple comparison** | | | | | |
|---|---|---|---|---|---|
| Number of families | | | | | 3 |
| Number of comparisons per family | | | | | 3 |
| Alpha | | | | | 0.05 |
| **Dunnett's multiple comparisons test** | **Mean Diff.** | **95.00% CI of diff.** | **Below threshold?** | **Summary** | **Adjusted P Value** |
| WT | | | | | |
| control vs. 20 µM Acon | -0.1233 | -0.6951 to 0.4485 | No | ns | 0.9072 |
| control vs. 50 µM Acon | -0.6267 | -1.198 to -0.05485 | Yes | * | 0.0296 |
| control vs. 100 µM Acon | 0.1667 | -0.4051 to 0.7385 | No | ns | 0.8072 |
| *sigRep* | | | | | |
| control vs. 20 µM Acon | -3.330 | -3.902 to -2.758 | Yes | **** | <0.0001 |
| control vs. 50 µM Acon | 1.510 | 0.9382 to 2.082 | Yes | **** | <0.0001 |
| control vs. 100 µM Acon | 2.450 | 1.878 to 3.022 | Yes | **** | <0.0001 |
| *gunSOS1* | | | | | |
| control vs. 20 µM Acon | -2.660 | -3.232 to -2.088 | Yes | **** | <0.0001 |
| control vs. 50 µM Acon | -0.1900 | -0.7618 to 0.3818 | No | ns | 0.7430 |
| control vs. 100 µM Acon | -0.2067 | -0.7785 to 0.3651 | No | ns | 0.6948 |

| **Test details** | **Mean 1** | **Mean 2** | **Mean Diff.** | **SE of diff.** | **N1** | **N2** | **q** | **DF** |
|---|---|---|---|---|---|---|---|---|
| WT | | | | | | | | |
| control vs. 20 µM Acon | 1.000 | 1.123 | -0.1233 | 0.2281 | 3 | 3 | 0.5407 | 24.00 |
| control vs. 50 µM Acon | 1.000 | 1.627 | -0.6267 | 0.2281 | 3 | 3 | 2.747 | 24.00 |
| control vs. 100 µM Acon | 1.000 | 0.8333 | 0.1667 | 0.2281 | 3 | 3 | 0.7307 | 24.00 |
| *sigRep* | | | | | | | | |
| control vs. 20 µM Acon | 3.130 | 6.460 | -3.330 | 0.2281 | 3 | 3 | 14.60 | 24.00 |
| control vs. 50 µM Acon | 3.130 | 1.620 | 1.510 | 0.2281 | 3 | 3 | 6.620 | 24.00 |
| control vs. 100 µM Acon | 3.130 | 0.6800 | 2.450 | 0.2281 | 3 | 3 | 10.74 | 24.00 |
| *gunSOS1* | | | | | | | | |
| control vs. 20 µM Acon | 0.2567 | 2.917 | -2.660 | 0.2281 | 3 | 3 | 11.66 | 24.00 |
| control vs. 50 µM Acon | 0.2567 | 0.4467 | -0.1900 | 0.2281 | 3 | 3 | 0.8330 | 24.00 |
| control vs. 100 µM Acon | 0.2567 | 0.4633 | -0.2067 | 0.2281 | 3 | 3 | 0.9060 | 24.00 |

**Supplementary Table 8. Statistical analyses for *PSBP2* expression following treatment with aconitate, presented in Fig. 6c.** *P*, probability (significance test); *SS*, sum of squares; *DF*, degree of freedom; *MS*, means square; *F*, F-value (Fisher test); *DFn*, degrees of freedom in the numerator; *DFd*, degrees of freedom in the denominator; *SE*, standard error of the sample mean; *N*, sample number.

| **Two-way ANOVA Ordinary** | | | | | |
|---|---|---|---|---|---|
| Alpha | 0.05 | | | | |
| **Source of Variation** | **% of total variation** | **P value** | **P value summary** | **Significant?** | |
| Interaction | 25.51 | <0.0001 | **** | Yes | |
| strain | 47.78 | <0.0001 | **** | Yes | |
| metabolite | 25.30 | <0.0001 | **** | Yes | |
| **ANOVA table** | **SS** | **DF** | **MS** | **F (DFn, DFd)** | **P value** |
| Interaction | 161.1 | 6 | 26.85 | F (6, 24) = 72.08 | P<0.0001 |
| strain | 301.7 | 2 | 150.9 | F (2, 24) = 405.0 | P<0.0001 |
| metabolite | 159.8 | 3 | 53.26 | F (3, 24) = 143.0 | P<0.0001 |
| Residual | 8.940 | 24 | 0.3725 | | |
| **Data summary** | | | | | |
| Number of columns (metabolite) | 4 | | | | |
| Number of rows (strain) | 3 | | | | |
| Number of values | 36 | | | | |

| **Two-way ANOVA Multiple comparison** | | | | | |
|---|---|---|---|---|---|
| Number of families | 3 | | | | |
| Number of comparisons per family | 3 | | | | |
| Alpha | 0.05 | | | | |
| **Dunnett's multiple comparisons test** | **Mean Diff.** | **95.00% CI of diff.** | **Below threshold?** | **Summary** | **Adjusted P Value** |
| WT | | | | | |
| control vs. 20 µM Acon | -0.3700 | -1.619 to 0.8792 | No | ns | |
| control vs. 50 µM Acon | -1.770 | -3.019 to -0.5208 | Yes | ** | |
| control vs. 100 µM Acon | -1.697 | -2.946 to -0.4474 | Yes | ** | |
| *sigRep* | | | | | |
| control vs. 20 µM Acon | -7.653 | -8.903 to -6.404 | Yes | **** | |
| control vs. 50 µM Acon | 0.3600 | -0.8892 to 1.609 | No | ns | |
| control vs. 100 µM Acon | 5.030 | 3.781 to 6.279 | Yes | **** | |
| *gunSOS1* | | | | | |
| control vs. 20 µM Acon | -5.497 | -6.746 to -4.247 | Yes | **** | |
| control vs. 50 µM Acon | -0.1767 | -1.426 to 1.073 | No | ns | |
| control vs. 100 µM Acon | -0.1600 | -1.409 to 1.089 | No | ns | |

| **Test details** | **Mean 1** | **Mean 2** | **Mean Diff.** | **SE of diff.** | **N1** | **N2** | **q** | **DF** |
|---|---|---|---|---|---|---|---|---|
| WT | | | | | | | | |
| control vs. 20 µM Acon | 1.000 | 1.370 | -0.3700 | 0.4983 | 3 | 3 | 0.7425 | 24.00 |
| control vs. 50 µM Acon | 1.000 | 2.770 | -1.770 | 0.4983 | 3 | 3 | 3.552 | 24.00 |
| control vs. 100 µM Acon | 1.000 | 2.697 | -1.697 | 0.4983 | 3 | 3 | 3.405 | 24.00 |
| *sigRep* | | | | | | | | |
| control vs. 20 µM Acon | 7.333 | 14.99 | -7.653 | 0.4983 | 3 | 3 | 15.36 | 24.00 |
| control vs. 50 µM Acon | 7.333 | 6.973 | 0.3600 | 0.4983 | 3 | 3 | 0.7224 | 24.00 |
| control vs. 100 µM Acon | 7.333 | 2.303 | 5.030 | 0.4983 | 3 | 3 | 10.09 | 24.00 |
| *gunSOS1* | | | | | | | | |
| control vs. 20 µM Acon | 0.1167 | 5.613 | -5.497 | 0.4983 | 3 | 3 | 11.03 | 24.00 |
| control vs. 50 µM Acon | 0.1167 | 0.2933 | -0.1767 | 0.4983 | 3 | 3 | 0.3545 | 24.00 |
| control vs. 100 µM Acon | 0.1167 | 0.2767 | -0.1600 | 0.4983 | 3 | 3 | 0.3211 | 24.00 |

**Supplementary Table 9. Statistical analyses for *MBS* expression following treatment with aconitate, presented in Fig. 6c.** *P*, probability (significance test); *SS*, sum of squares; *DF*, degree of freedom; *MS*, means square; *F*, F-value (Fisher test); *DFn*, degrees of freedom in the numerator; *DFd*, degrees of freedom in the denominator; *SE*, standard error of the sample mean; *N*, sample number.

| **Two-way ANOVA Ordinary** | | | | | |
|---|---|---|---|---|---|
| Alpha | 0.05 | | | | |
| **Source of Variation** | **% of total variation** | **P value** | **P value summary** | **Significant?** | |
| Interaction | 28.05 | <0.0001 | **** | Yes | |
| strain | 32.20 | <0.0001 | **** | Yes | |
| metabolite | 37.62 | <0.0001 | **** | Yes | |
| **ANOVA table** | **SS** | **DF** | **MS** | **F (DFn, DFd)** | **P value** |
| Interaction | 1424 | 6 | 237.3 | F (6, 24) = 52.66 | P<0.0001 |
| strain | 1635 | 2 | 817.4 | F (2, 24) = 181.4 | P<0.0001 |
| metabolite | 1910 | 3 | 636.5 | F (3, 24) = 141.2 | P<0.0001 |
| Residual | 108.2 | 24 | 4.507 | | |

| **Data summary** | |
|---|---|
| Number of columns (metabolite) | 4 |
| Number of rows (strain) | 3 |
| Number of values | 36 |

| **Two-way ANOVA Multiple comparison** | |
|---|---|
| Number of families | 3 |
| Number of comparisons per family | 3 |
| Alpha | 0.05 |

| **Dunnett's multiple comparisons test** | **Mean Diff.** | **95.00% CI of diff.** | **Below threshold?** | **Summary** | **Adjusted P Value** |
|---|---|---|---|---|---|
| WT | | | | | |
| control vs. 20 µM Acon | -0.4133 | -4.759 to 3.932 | No | ns | 0.9904 |
| control vs. 50 µM Acon | -4.537 | -8.882 to -0.1912 | Yes | * | 0.0394 |
| control vs. 100 µM Acon | -3.460 | -7.806 to 0.8855 | No | ns | 0.1395 |
| *sigRep* | | | | | |
| control vs. 20 µM Acon | -25.88 | -30.23 to -21.54 | Yes | **** | <0.0001 |
| control vs. 50 µM Acon | -14.28 | -18.63 to -9.934 | Yes | **** | <0.0001 |
| control vs. 100 µM Acon | 1.287 | -3.059 to 5.632 | No | ns | 0.8002 |
| *gunSOS1* | | | | | |
| control vs. 20 µM Acon | -28.43 | -32.78 to -24.08 | Yes | **** | <0.0001 |
| control vs. 50 µM Acon | -0.2200 | -4.566 to 4.126 | No | ns | 0.9985 |
| control vs. 100 µM Acon | -0.3267 | -4.672 to 4.019 | No | ns | 0.9952 |

| **Test details** | **Mean 1** | **Mean 2** | **Mean Diff.** | **SE of diff.** | **N1** | **N2** | **q** | **DF** |
|---|---|---|---|---|---|---|---|---|
| WT | | | | | | | | |
| control vs. 20 µM Acon | 1.000 | 1.413 | -0.4133 | 1.733 | 3 | 3 | 0.2384 | 24.00 |
| control vs. 50 µM Acon | 1.000 | 5.537 | -4.537 | 1.733 | 3 | 3 | 2.617 | 24.00 |
| control vs. 100 µM Acon | 1.000 | 4.460 | -3.460 | 1.733 | 3 | 3 | 1.996 | 24.00 |
| *sigRep* | | | | | | | | |
| control vs. 20 µM Acon | 9.413 | 35.30 | -25.88 | 1.733 | 3 | 3 | 14.93 | 24.00 |
| control vs. 50 µM Acon | 9.413 | 23.69 | -14.28 | 1.733 | 3 | 3 | 8.238 | 24.00 |
| control vs. 100 µM Acon | 9.413 | 8.127 | 1.287 | 1.733 | 3 | 3 | 0.7423 | 24.00 |
| *gunSOS1* | | | | | | | | |
| control vs. 20 µM Acon | 0.4633 | 28.89 | -28.43 | 1.733 | 3 | 3 | 16.40 | 24.00 |
| control vs. 50 µM Acon | 0.4633 | 0.6833 | -0.2200 | 1.733 | 3 | 3 | 0.1269 | 24.00 |
| control vs. 100 µM Acon | 0.4633 | 0.7900 | -0.3267 | 1.733 | 3 | 3 | 0.1884 | 24.00 |

**Supplementary Table 10. Statistical analyses for *SAK1* expression following treatment with aconitate, presented in Fig. 6c.** *P*, probability (significance test); *SS*, sum of squares; *DF*, degree of freedom; *MS*, means square; *F*, F-value (Fisher test); *DFn*, degrees of freedom in the numerator; *DFd*, degrees of freedom in the denominator; *SE*, standard error of the sample mean; *N*, sample number.

| **Two-way ANOVA Ordinary** | | | | | |
|---|---|---|---|---|---|
| Alpha | 0.05 | | | | |
| **Source of Variation** | **% of total variation** | **P value** | **P value summary** | **Significant?** | |
| Interaction | 49.81 | <0.0001 | **** | Yes | |
| strain | 12.86 | <0.0001 | **** | Yes | |
| metabolite | 35.02 | <0.0001 | **** | Yes | |
| **ANOVA table** | **SS** | **DF** | **MS** | **F (DFn, DFd)** | **P value** |
| Interaction | 15402 | 6 | 2567 | F (6, 24) = 86.01 | P<0.0001 |
| strain | 3976 | 2 | 1988 | F (2, 24) = 66.62 | P<0.0001 |
| metabolite | 10830 | 3 | 3610 | F (3, 24) = 121.0 | P<0.0001 |
| Residual | 716.3 | 24 | 29.84 | | |

| **Data summary** | |
|---|---|
| Number of columns (metabolite) | 4 |
| Number of rows (strain) | 3 |
| Number of values | 36 |

| **Two-way ANOVA Multiple comparison** | |
|---|---|
| Number of families | 3 |
| Number of comparisons per family | 3 |
| Alpha | 0.05 |

| **Dunnett's multiple comparisons test** | **Mean Diff.** | **95.00% CI of diff.** | **Below threshold?** | **Summary** | **Adjusted P Value** |
|---|---|---|---|---|---|
| WT | | | | | |
| control vs. 20 µM Acon | -0.2367 | -11.42 to 10.95 | No | ns | >0.9999 |
| control vs. 50 µM Acon | -3.273 | -14.46 to 7.909 | No | ns | 0.8052 |
| control vs. 100 µM Acon | -2.583 | -13.77 to 8.599 | No | ns | 0.8895 |
| *sigRep* | | | | | |
| control vs. 20 µM Acon | -16.06 | -27.25 to -4.881 | Yes | ** | 0.0040 |
| control vs. 50 µM Acon | -5.853 | -17.04 to 5.329 | No | ns | 0.4296 |
| control vs. 100 µM Acon | 2.253 | -8.929 to 13.44 | No | ns | 0.9222 |
| *gunSOS1* | | | | | |
| control vs. 20 µM Acon | -107.2 | -118.4 to -96.01 | Yes | **** | <0.0001 |
| control vs. 50 µM Acon | -0.9400 | -12.12 to 10.24 | No | ns | 0.9933 |
| control vs. 100 µM Acon | -0.6300 | -11.81 to 10.55 | No | ns | 0.9979 |

| **Test details** | **Mean 1** | **Mean 2** | **Mean Diff.** | **SE of diff.** | **N1** | **N2** | **q** | **DF** |
|---|---|---|---|---|---|---|---|---|
| WT | | | | | | | | |
| control vs. 20 µM Acon | 1.000 | 1.237 | -0.2367 | 4.461 | 3 | 3 | 0.05306 | 24.00 |
| control vs. 50 µM Acon | 1.000 | 4.273 | -3.273 | 4.461 | 3 | 3 | 0.7338 | 24.00 |
| control vs. 100 µM Acon | 1.000 | 3.583 | -2.583 | 4.461 | 3 | 3 | 0.5792 | 24.00 |
| *sigRep* | | | | | | | | |
| control vs. 20 µM Acon | 7.743 | 23.81 | -16.06 | 4.461 | 3 | 3 | 3.601 | 24.00 |
| control vs. 50 µM Acon | 7.743 | 13.60 | -5.853 | 4.461 | 3 | 3 | 1.312 | 24.00 |
| control vs. 100 µM Acon | 7.743 | 5.490 | 2.253 | 4.461 | 3 | 3 | 0.5052 | 24.00 |
| *gunSOS1* | | | | | | | | |
| control vs. 20 µM Acon | 0.8933 | 108.1 | -107.2 | 4.461 | 3 | 3 | 24.03 | 24.00 |
| control vs. 50 µM Acon | 0.8933 | 1.833 | -0.9400 | 4.461 | 3 | 3 | 0.2107 | 24.00 |
| control vs. 100 µM Acon | 0.8933 | 1.523 | -0.6300 | 4.461 | 3 | 3 | 0.1412 | 24.00 |

**Supplementary Table 11. Primers used in this study.**

| v5.5 gene ID | Gene name | Forward (5' → 3') | Reverse (5' → 3') |
|---|---|---|---|
| ***GPX5-ARS2* fusion** | | | |
| Cre10.g458450 | *GPX5 5'RR* | CTCGAGGGTACATGTTTAGAACCCGCT* | GATATCTGCAATCGTCGCTGGTTC* |
| **Rescue with genomic *TSPP1* (BAC PTQ5987 template)** | | | |
| Cre12.g497750 | *TSPP1* | GATGAAGCTCAGGGCGAGAC | CAGCAGCAGTAGCGAACCAA |
| **qRT-PCR** | | | |
| Reference gene for all qRT-PCR | *18S rRNA* | GATGGCTACCACATCCAAGGAA[a] | AAGCGCCCGGTATTGTTATTTATT[a] |
| Cre10.g458450 | *GPX5*$_{cyt}$ | GCGGTCGCCAATAACCAAT[a] | AAGGGCTGTCCCGAAAGC[a] |
| Cre10.g458450 | *GPX5*$_{cp}$ | AACCCTTTCACTCACATGCTGTCT[a] | CGAGCGGCGACAGGAGTA[a] |
| | *GPX5-ARS2* | GCCTTTGCATCTACTGAACCAG | TCCCAAACGATCCCTTGACAG |
| Cre12.g497750 | *TSPP1* | GAAGATAGCAATGACGGCAAGG | ATGCTTCGTCCTTCTCTGACTG |
| Cre12.g497750 | *TSPP1* (kinetics) | AGGACGAAGCATACCTCAAGTG | TGAGCGTGCCATCATAGTCTAG |
| Cre16.g678851 | *PSBP2* | AAGCTGTACGAGTACGAGTACG | CTTGTATGCCGCGTTTAGGATG |
| Cre09.g416500 | *MBS* | TGCGCGGATCTACCAAGAAG | CGGAAGCGAAGTGTACATCTTC |
| Cre17.g741300 | *SAK1* | TCAAGCGTGTGGGTAAGAGCTA[b] | ACGCTATCTCCGTCCTAATCCA[b] |
| Cre06.g281250 | *CFA1* | CCTACAACGACAACGACGTG[b] | GGAAGTTCCAGGATGACCAG[b] |
| Cre09.g398700 | *CFA2* (*CPLD27*) | GTCCATTGAGATGTTCGAGCAC | ATGTGCACGAACAACTTGCC |
| Cre06.g299700 | *SOUL1* | TGAAGAAGATCCCCATGACTGC | CGAAGAACGACACCTTGAAGTG |
| Cre08.g380300 | *MSRA3* | GACTGAAGTTGGCGACGTTC | AGTCGTAGTTGGGGTTCTTGTC |
| Cre14.g623650 | *ADH7* | AAGAAGGTCGTTGGCTCCATC | TAGCCTCGTTCACCTTGCTG |
| Cre12.g503950 | *RABPR2* | AGGACTACAGCAAGCAGATCG | TACCAGCGCAGGTAGTTGTC |
| Cre01.g007300 | | TAGGGCCAAAGTTCTGTAACCC | GTGGCCGTCAGTGTTAATCTTG |
| Cre16.g676150 | *MSD3* | TCGACAACGAGACCATGTTCC | ACAATCTCCGACAGCGACAG |
| Cre13.g588150 | *VTC2* | CAACCAGCCGTTCAACATCATC | CCGCAATCTCAAACGATGCC |
| Cre10.g456750 | *DHAR1* | GACAGCGATGTCATTGTGGTG | AACAGTTTGGCGCCGATTTC |
| Cre16.g688550 | *GSTS1* | TGTTGTTCCACATCGGCAAC | CCTGGCCAAATGGGAACTTTG |
| Cre10.g436050 | *FSD1* | TTCTTCTGGGAGAGCATGAAGC | ACTCCTCCTTGAACTTGTCCAG |
| Cre01.g020223 | *FUM2* | GCAGATGAGTTTGCAGGCATC | TACTTCACCTGCGCTGCATAG |
| Cre17.g732802 | | ATCCAGAAGTGGGAGTACGTG | ATCCAACGTCACCACCCATG |
| Cre01.g042750 | *ACH1* | CCATGTTCCCTTACAACAAGCG | TTCAGGTGCTCCTTGAACGAG |

*Underlined sequences indicate restriction sites, XhoI (CTCGAG) and EcoRV (GATATC); [a]Fischer et al. (2009); [b]Wakao et al. (2014)
